# Inhibition of a Selective SWI/SNF Function Synergizes with ATR Inhibitors in Cancer Cell Killing

**DOI:** 10.1101/660456

**Authors:** Emma J. Chory, Jacob G. Kirkland, Chiung-Ying Chang, Vincent D. D’Andrea, Sai Gourinsankar, Emily C. Dykhuizen, Gerald R. Crabtree

## Abstract

SWI/SNF (BAF) complexes are a diverse family of ATP-dependent chromatin remodelers produced by combinatorial assembly that are mutated in and thought to contribute to 20% of human cancers and a large number of neurologic diseases. The gene-activating functions of BAF complexes are essential for viability of many cell types, limiting the development of small molecule inhibitors. To circumvent the potential toxicity of SWI/SNF inhibition, we identified small molecules that inhibit the specific repressive function of these complexes but are relatively non-toxic and importantly synergize with ATR inhibitors in killing cancer cells. Our studies suggest an avenue for therapeutic enhancement of ATR/ATM inhibition and provide evidence for chemical synthetic lethality of BAF complexes as a therapeutic strategy in cancer.

## INTRODUCTION

The accessibility of the genome to DNA repair, recombination, and transcriptional machinery is regulated by several processes including DNA methylation, histone modifications and ATP-dependent chromatin remodeling. The human genome encodes 29 ATPases related to the chromatin remodeling ATPase, SWI2, which was first discovered in yeast^1,2^. These ATPases are largely non-redundant and often observed as multi-subunit assemblies. Among this family of ATP-dependent chromatin regulators, mSWI/SNF or BAF (Brg Associated Factor) complexes are highly mutated in human cancer^3^ and appear to play dose-dependent roles in tumor suppression^4^, neural development^5^, oncogenesis^6^, and maintaining HIV latency^7–9^. These complexes are combinatorially assembled from 16 subunits encoded by 31 genes **(Figure 1A)** giving them specific roles in many biologic processes. Recently, BAF complexes have been found to take on non-canonical assemblies (ncBAF) ^10,11,58^, which are essential in malignant rhabdoid tumors^10^ and synovial sarcoma^11^ **(Figure 1A)**. Although many of the subunits are required for essential cellular mechanisms^12,13^, other subunits have highly selective functions^14–17^. For example, the nBAF complex, found only in neurons, is also combinatorially assembled and plays instructive roles in reprograming fibroblasts to neurons^17^. The highly selective nature of these protein assemblies raises the possibility to develop specific inhibitors with precise therapeutic roles. Despite being mutated in roughly 20% of human cancers^18^, there is a significant dearth of SWI/SNF inhibitors with utility in treating cancer. For example, PFI-3, which targets the bromodomain of ATPase containing subunit^19^, has been shown to have no measurable effects on inhibiting the growth of cancer cells^20^, despite its ability to impair trophoblast development. A class of phospho-aminoglycosides (phospho-kanamycin) inhibits the yeast SWI2/SNF2 complex but also have limited utility in mammalian cells, are relatively non-specific ATPase inhibitors, and would likely be highly toxic in this context^21^. Recently, a BRG1 ATPase inhibitor was discovered^22^, yet its non-specific inhibition of both BRG1 and BRM would likely be toxic in patients due to the essential nature of the core ATPase subunits in cellular viability in many cell types.^10^

**Figure 1:**
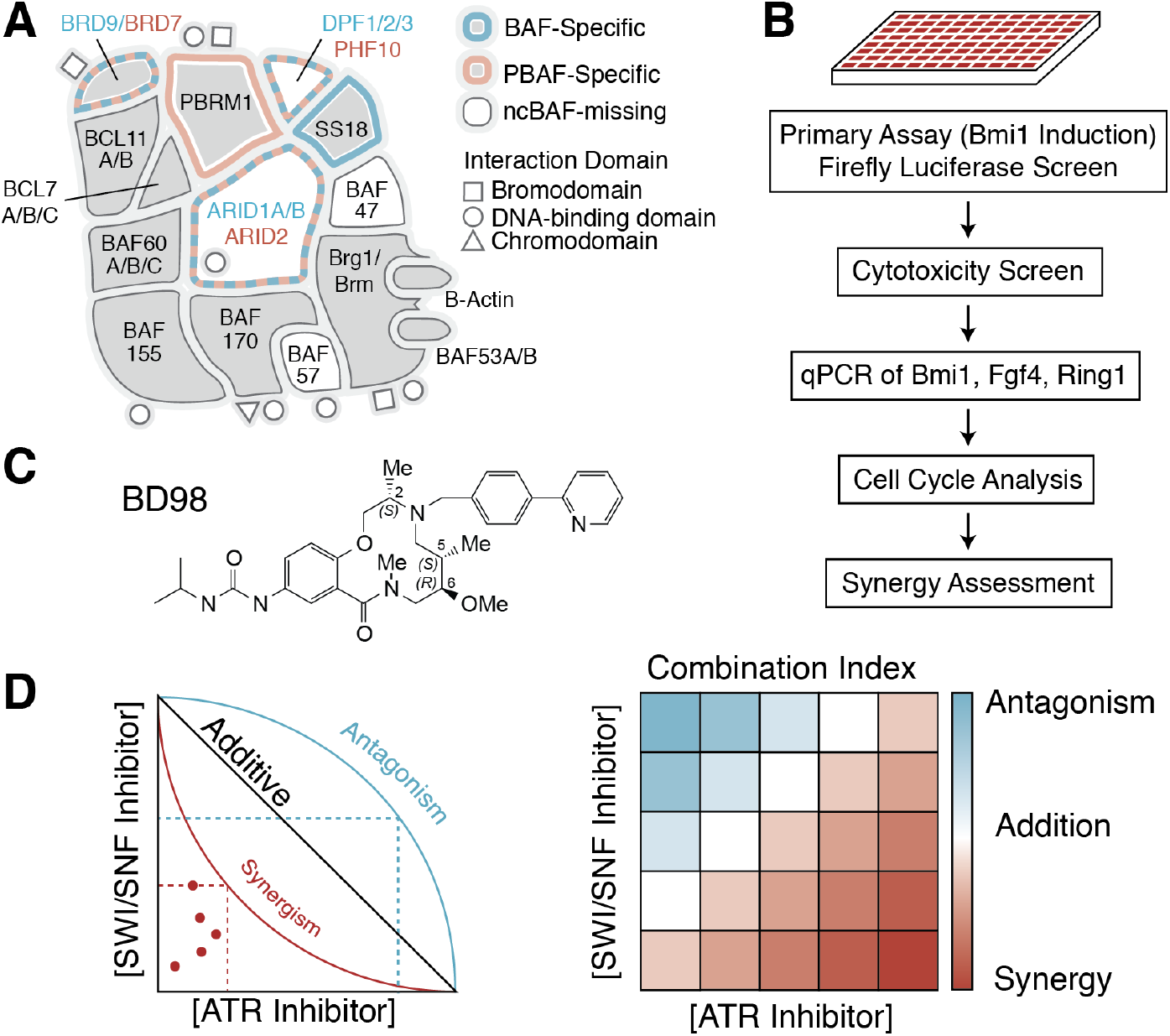
Strategy for assessing hyper-synthetic lethality of combination BD98 and ATRi. **(A)** Macromolecular assembly of SWI/SNF complexes. BAF/PBAF-specific subunits labelled in blue/red respectively. **(B)** Workflow of screen used to identify putative BAF inhibitors. **(C)** Structure of BD98. (**D)** Summary of results (combination index scores, normalized isobolograms) obtained in Chou-Talalay method for assessment of synergy

As members of the thrithorax group proteins, mSWI/SNF complexes are generally thought to be activators of transcription. However, they can also directly inhibit expression of PRC1 genes in murine embryonic stem cells^23^. To exploit the repressive functions of BAF complexes, we previously developed screens for one of these specialized functions: the ability to repress the *Bmi1* gene^24^. Utilizing a luciferase reporter gene inserted in-frame into the *Bmi1* locus, we screened for molecules that could specifically de-repress its expression in mouse embryonic stem cells (mESCs) **(Figure 1B)**^24^. This strategy had the particular advantage of being able to selectively screen for BAF functionality orthogonal to the ATPase function, which would likely result in high levels of toxicity or even oncogenesis. We identified multiple 12-membered macrolactam candidates from a library of diversity-oriented-synthesis molecules^25,26^ **(Figure 1C)** which appear to bind to ARID1A-specific complexes^27^. These inhibitors are remarkably non-toxic in most cell lines and do not appear to independently produce the full decatenating checkpoint arrest found with deletion or depletion of essential subunits such as BRG1 (SMARCA4), BAF57 (SMARCE1) or BAF53a (ACT6LA)^27^. This class of structures appears to bind to complexes containing homologous BAF250a/b (ARID1A/B) subunits^27^ which play critical, but redundant roles in mammalian BAF function, and are specific to canonical-BAF complexes^28^ **(Figure 1A)**. However, it is likely that the molecules bind an interfacial surface of BAF250 produced by the junction with another subunit which has made precise target-identification of the binding site non-trivial.

The decatenation checkpoint arrest observed upon deletion of essential subunits of the BAF complex appears to be related to the inability of Topoisomerase IIa to bind to DNA and relieve catenated chromosomes at mitosis^13^. This results in anaphase bridges and cell cycle arrest mediated by ATR kinase (Ataxia-Telangiectasia Mutated and Rad3-related protein kinase) ^29,30^. Having recently validated this macrolactam class of SWI/SNF inhibitory molecules as reversal agents for HIV latency^27,31^, we sought to ascertain whether these inhibitors of the repressive function of BAF may provide a therapeutic strategy for cancer, without resulting in broad toxicity, as would be expected from BRG1 ATPase inhibitors. Recently, inhibitors of the ATR kinase demonstrated enhanced chemotoxic effects of DNA damaging agents^32,33^. ATR and ATM are known to phosphorylate BAF170 (SMARCC2) and BRG1 and induce the localization of these complexes to sites of DNA damage^15^. In addition, potent small molecule inhibitors of ATR have been shown to induce a synthetic lethal function in cancer cell lines depleted of the ARID1A/B subunits of the BAF complex^27^. This synthetic lethal interaction has been attributed to the increased dependency of ARID1A/B-deficient cells on the ATR-mediated G2/M decatenation checkpoint following loss of ARID1A/B-mediated interactions of the BAF complex with TOP2A^13,34^. These results suggested that ATR and SWI/SNF inhibition might functionally synergize because they work on contingent steps^34^. To address the potential of BAF-inhibitory molecules to function as chemical-genetic mimetics, we developed a high-throughput Chou-Talalay method of synergy assessment (**Figure 1D**) (Chory & Divakaran, unpublished), to demonstrate a synergistic, chemical-synthetic lethality in cancers containing canonical BAF complexes, when combining SWI/SNF and ATR inhibitors. Our data indicate that the use of selective, non-toxic BAF inhibitors might provide a strategy for enhancement of chemotherapeutics in cancer.

## RESULTS AND DISCUSSION

### SWI/SNF inhibition sensitizes cancer cells to ATR inhibition

We recently reported a medium-throughput screen to identify small molecules that result in increased expression of *Bmi1*, a known genetic target of BAF whose expression is increased about 10-fold upon deletion of the *Smarca4* subunit of the BAF complex^26^. Through the development of a high-throughput, *Bmi1* luciferase reporter assay, we identified multiple 12-membered macrolactam candidates from a library of diversity-oriented-synthesis^25^ molecules which exhibited stereospecificity and provided insight into a structure-activity-relationship^27^. While many of the molecules we identified blocked cell cycle progression and had toxic effects (as might be expected from a general inhibitor of the BAF complex), one class of molecules was both non-toxic and exhibited robust activation of *Bmi1* as well as other BAF targets, suggesting that it selectively inhibited the repressive functions of the canonical BAF complex. Of the most potent molecules, BRD-K98645985 (which we refer to as BD98) (**Figure 1C**) was selected for further analysis for synergy with the ATR inhibitor VE-821 on the viability of cancer cells containing intact BAF complexes. We have previously shown that BD98 is able to selectively bind and *in vitro* thermo-stabilize ARID1A/B (BAF250A/B)^27^, enrich BAF subunits ARID1A, BAF155, PBRM1, and BAF47 by small-molecule pull-down, and activate *Bmi1* at an EC_50_ of ~2.4 uM^27^. Hence, we loosely refer to it as a BAF inhibitor although its direct binding target has not been identified with complete confidence. Previous studies showed that upon acute deletion of *ARID1A*, the IC50 of the ATR inhibitor VE-821 in HCT116 colorectal cancer cells shifted from ~10uM to ~1uM after 5 days of treatment^34^. In this study, we treated HCT116 cells with increasing doses of VE-821 (1uM-50uM) and increasing doses of BD98 (1uM–30uM) independently for 5 days to establish the respective dose responses of the two **(Figure 2A)**. Then, cells were treated with all possible combinations of 5 doses of both VE-821 (1.25-20uM) BD98 (1.25-20uM) for a total of 25 total combinations, in 8 replicates (**Figure 2B**). By simultaneously treating HCT116 cells with increasing concentrations of VE-821 and the putative BAF inhibitor, BD98, we observed dose-responsive decreases in VE-821 IC50s ranging from ~4uM at the lowest dose of BD98 (1.25uM) to an IC50 ~1uM at the highest dose tested in combination (**Figure 2B**). This indicated that co-inhibition of both BAF and ATR could phenocopy the dose-responsive shift previously observed by ARID1A knockdown in HCT116 cells ^34^.

**Figure 2:**
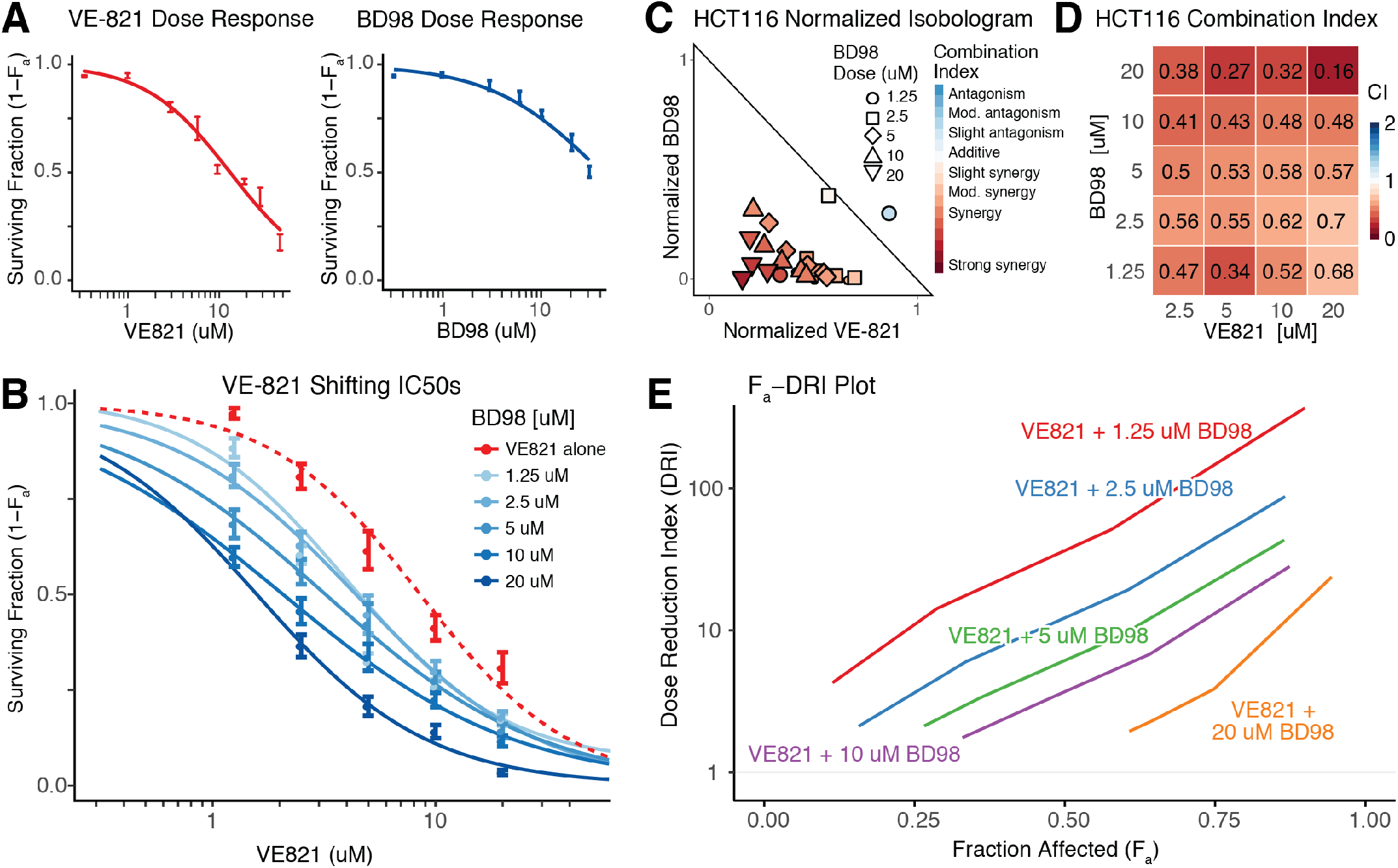
BAF and ATR Inhibition is synergistic. **(A)** Dose response curves of HCT116 cell line exposed to increasing concentrations of ATR inhibitor (VE-821) and putative BAF inhibitor (BD98) for 5 days. **(B)** Shifting IC50 dose-response curves of HCT116 cells treated with VE-821 and increasing doses of BD98 for 5 days. **(C)** Normalized isobologram of synergy between VE-821 and BD98 in HCT116 cells. **(D)** Combination index values quantify synergy between BD98 and VE-821. **(E)** Plot of the dose reduction index of increasing concentrations of VE-821 and 5 concentrations of BD98 against the fraction of affected cells.

### Combination BD98/ATRi treatment is synergistic

Occasionally, combining two drugs that act on contingent steps will result in enhanced potency. To assess whether the combined effect is greater than the predicted individual potencies, and hence truly synergistic, requires a rigorous quantitative analysis of both the independent and combined effects of the two molecules, through the generation of isobolograms and quantification of a combination index (**Figure 1D**). Synergism in drug combinations allows for the use of lower doses of both molecules, which can reduce adverse effects. We first aimed to validate the combination of VE-821 and BD98 for synergistic therapeutic value. To assess drug synergy, the Chou-Talalay method was employed to provide a mechanism-independent method to quantify the synergism of drug interactions^35^. This method combines elements of the Scatchard, Michaelis-Menten, Hill, and Henderson-Hasslebalch equations through the law of mass-action^36,37^ to generate a statistical and quantifiable assessment of synergy. Molecules were tested both independently and then in 25 different combinations in HCT116 cells for 5 days as described above. A combination index (CI), which is a statistically-significant representation of synergy, was determined by generating median-effect equations of the two-respective from their respective dose responses. The median-effect equation was used to calculate a median-effect dose, and (D_x_)_ATRi,BD98_ values which correspond for the respective doses of the BD98 and ATRi compounds and correlate with a given percentage of cells affected by the individual treatments. Normalized index values (I_ATRi_, I_BD98_), and the combination index are a ratio between the treatment dose, and the D_x_ for a given fraction of affected cells. Combination indices less than 0.9 are statistically synergistic, while values ranging from 0.9 to 1.1 are additive, and CIs above 1.1 are antagonistic. Isobolograms and combination index values were calculated for the tested dose combinations of VE-821 and BD98 (**Figure 2C-2D**). The average combination index observed in HCT116 cells was 0.53 ± 0.05 which indicates clear synergism between VE-821 and BD98 (**Figure 2D**). Further, the dose reduction index (an inverse relation to the combination index) was calculated to demonstrate that at low concentrations, the combination of VE-821 and BD98 can reduce the dosing over 10-fold, while at higher concentrations of VE-821 the dose reduction index reaches nearly 100 (**Figure 2E**), indicating that the combination has the potential to significantly lower the dosing of the toxic ATR inhibitor.

### Putative SWI/SNF inhibitor phenocopies ARID1A/B loss

To analyze the effects of ARID1A on the synergistic combination, we made use of HCT116 cells for which we have both isogenic *ARID1A*^+/+^ and *ARID1A*^−/−^ cell lines. The primary target of our original screen (*Bmi1*), is induced by loss of SMARCA4, which is present in human BAF, PBAF, and ncBAF complexes ^3,28^ **(Figure 1A)**. PBAF complexes differ from BAF complexes by incorporation of orthogonal subunits, specifically, ARID2 (as opposed to ARID1A/B), PHF10 (BAF45a) (as opposed to DPF10/BAF45d), and PBRM1. PBAF-specific subunits are also frequently mutated in cancer, specifically renal clear cell carcinomas and cholangio-carcinomas^38–41^. To narrow whether in HCT116 cells, the lead compound is acting on BAF or PBAF-related pathways, we constitutively knocked down ARID1A or ARID2 by lentiviral transduction of *ARID1A*^+/+^ HCT116 cells and compared the BD98/ATRi synergy to the wild-type and knockout (*ARID1A*^+/+^, *ARID1A*^−/−^) cells. Knockdown shARID1A cells demonstrated increased sensitivity to VE-821 as expected, as did the homozygous loss-of-function *ARID1A* HCT116 (*ARID1A*^−/−^) cell line, which contains mutations p.Q456*/p.Q456*. In contrast, shARID2 cells responded to ATR treatment similar to wildtype cells (**Figure 3A**). Importantly, upon knockdown of ARID1A or ARID2 (**Figure 3B**), the synergistic effect observed between BD98 and VE-821 was only ablated upon loss of ARID1A (**Figure 3C-E**) (CI_shARID1A_ = 1.48 ± 0.14, CI_ARID1A(−/−)_ = 1.44 ± 0.12), but not loss of shARID2 (CI_shARID2_ = 0.42 ± 0.04) (**Figure 3D-E**). This suggests that in the context of colorectal cancer, BD98 is acting through ARID1A-dependent mechanisms, or specific sub-units and/or functions of such complexes. Additionally, both shARID1A and *ARID1A*^−/−^ cells were slightly, but not-significantly less sensitive to BD98 than wild-type HCT116 cells. This minor effect is likely due to the fact that ARID1A is partially redundant with ARID1B in colon cancer, but could also be related to an off-target effect independent of ARID1 proteins.

**Figure 3:**
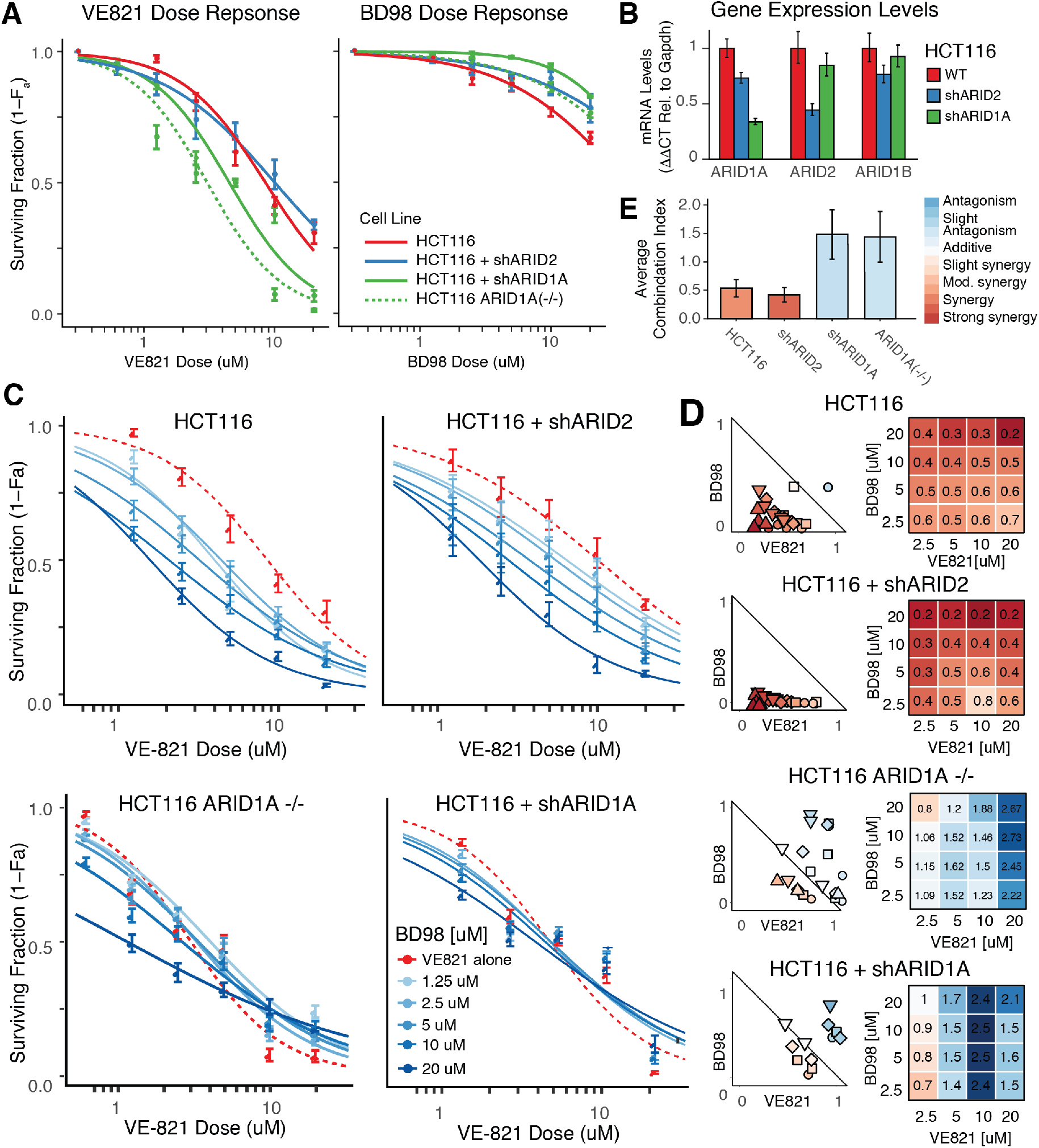
Putative BAF Inhibitor, BD98 phenocopies knockdown of ARID1A. **(A)** Cell survival data from HCT116 cells infected with shRNA lentivirus targeting ARID2 (blue), ARID1A (green), Control (red), and HCT116(ARID1A −/−) cells (green-dashed). Following lentiviral transduction and selection, cells were exposed to VE-821 or BD98 for 5 continuous days. Error bars represent s.d. of eight technical replicates in three separate experiments in 384-well plates. **(B)** Gene expression of ARID1A, ARID2, and ARID1B in HCT116 (WT, shARID1A, shARID2). Delta-Delta CT values compared to Gapdh and WT HCT116. **(C)** Shifting IC50 plot of WT HCT116 cells, ARID1A −/− HCT116 cells, and HCT116 cells infected with shRNA lentivirus targeting ARID2 or ARID1A treated with VE-821 and increasing doses of BD98 for 5 days. **(D)** Normalized isobologram plots of WT HCT116 cells, ARID1A −/− HCT116 cells, and HCT116 cells infected with shRNA lentivirus targeting ARID2 or ARID1A (as treated in 3C). **(E)** Average combination indexes for HCT116 cells infected with shRNA lentivirus targeting ARID2, ARID1A, WT, and HCT116 (ARID1A−/−).

### Analysis of additional library screen hits

From our original *Bmi1*-luciferase screen^26^, we chose 8 representative compounds to compare to the lead candidate which were either commercially available, or structurally similar to the 12-member macrocyclic lactam stereoisomer backbone of BD98. We have published an analysis of striking stereospecificity of the BD98 related molecules^27^, so we sought to assess whether any additional molecules from our primary screen would phenocopy ATRi sensitivity. First, we tested their ability to arrest mouse embryonic stem cells (mESCs) in G2/M but observed that G2/M arrest in mESCs was not directly correlated with the *Bmi1* and *Ring1a* induction observed in the primary screen (**Figure 4A-4B**). This suggests that many initial hits may be disrupting different SWI/SNF related mechanisms^13^, or may be non-specific *Bmi1* activators. Next, synergy was assessed between each hit and VE-821, as described above. We observed that the two structurally similar molecules (2A03 and 2C05), indeed demonstrated comparable synergy with VE-821 in HCT116 cells (CI_2A03_ = 0.65±0.05, CI_2C05_ = 0.69±0.08) (**Figure 4C-D**). However, the administration of all other compounds tested resulted in strictly additive or antagonistic effects when combined with VE-821 (**Supplemental Figure 1A**). Interestingly, the published PFI-3 BRG1 bromodomain inhibitor showed no measurable dose-responsive or synergy effects in the *ARID1A*^+/+^ or *ARID1A*^−/−^ HCT116 cell lines (**Supplemental Figure 1B**), suggesting that inhibition of the acetyl-lysine-binding function of BRG1 is not the major mechanism through which synergy is observed. These studies underly the specificity of the action of the structurally related SWI/SNF inhibitors compared to other hits from our *Bmi1*-induction screen, that also derepress the HIV LTR and terminate HIV latency^24,27^.

**Figure 4:**
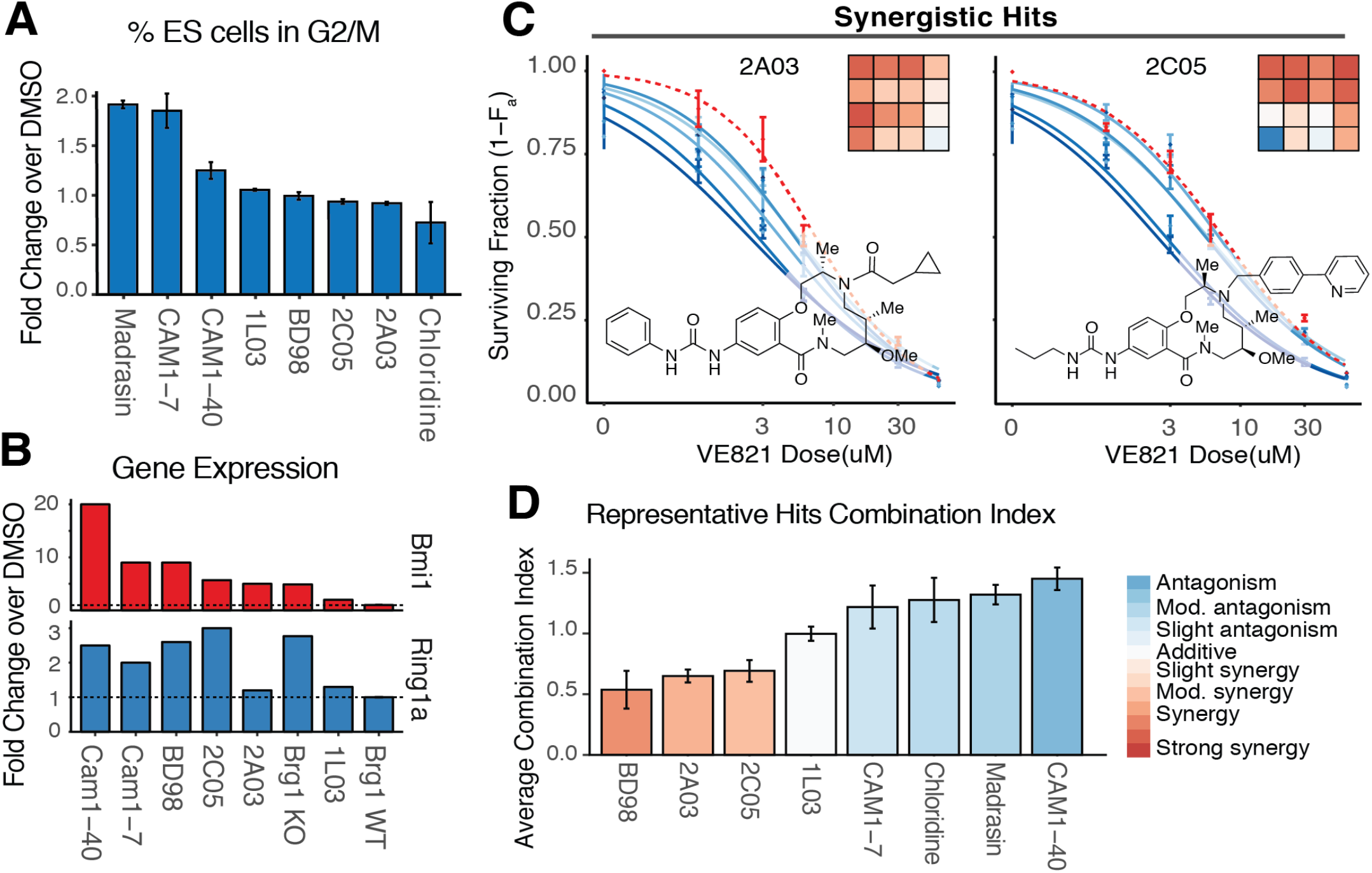
Structure Activity Relationship of Putative BAF Inhibitors. **(A)** Assessment of toxicity of putative inhibitors in ARID1a+/+, ARID1A−/− HCT116 cells and shARID1A ARID1a+/+ HCT116 cells by cell cycle analysis arrest in ESCs after release from double thymidine block. Data represent three separate cell-cycle analyses. **(B)** Induction of Bmi1 and Ring1a expression of 6 putative hits, compared to WT mESCs or Brg1 f/f actin-CreER mESCs following tamoxifen treatment. Data represents results from original screen. **(C)** Structures, shifting dose response, and CI grid of putative inhibitors demonstrating synergism: 2A03 and 2C05 following treatment with increasing doses of VE-821 (1-30uM) and increasing doses of each putative inhibitor (1-30uM) for 5 days. **(D)** Average combination indexes for HCT116 treated with putative BAF inhibitors and VE-821 (as treated in 4C).

### ATRi/BD98 Inhibition results in cell cycle defects

Given that we find that BD98 is largely non-toxic in mESCs, we sought to understand the mechanism by which BD98 kills cells in the presence of ATR inhibitors. To investigate this, we synchronized cells in the early G1/S phase and then tracked their progression through the cell cycle upon release into media containing BD98. To assess the full cell cycle, time points were collected beginning at early G0/G1 of the second cell division following release from thymidine block. We observed that HCT116 *ARID1A*^−/−^ cells exhibited slightly delayed progression through S-Phase compared to both acutely BD98 treated (10μM) and untreated HCT116 *ARID1A*^+/+^ cells, but overall progressed through the cell cycle at similar rates (~11hr per cycle) (**Figure 5A-5B)**. This is consistent with our observation that BD98 independently has minimal toxicity or effects on proliferation. We were curious as to the mechanism by which BD98 in combination with ATRi kills cells, so we further tracked the progression of cells released into media containing VE-821 (10μM) in the presence or absence of BD98 and compared to *ARID1A*^−/−^ cells under the same conditions (**Figure 5B**). Consistent with previous results^34^, we observed that HCT116 *ARID1A*^−/−^ cells displayed delayed progression through S phase, which is exacerbated by ATRi through relief of the pile-up and premature entry into mitosis (**Figure 5A-5B**). In addition, when treated with 10uM BD98 in combination with VE-821, WT HCT116 cells similarly exhibited delayed progression through S Phase (**Figure 5B-C**), comparable to ARID1A knockout. This result suggests that delayed progression through S Phase as a result of BD98, is likely due to either an inability of ARID1A-containing complexes to repair DNA damage, or perhaps collapsed/stalled replication forks, which result from defective function of the BAF complex^42^. We have previously demonstrated that deletion of *Brg1* in mouse embryonic stem cells (mESCs) results in the appearance DNA bridges during anaphase^13^. To examine whether inhibiting loss of BAF complexes results in increased double stranded breaks, we turned to mES cells which have stable genomes, in contrast to HCT116 cells. Using *Brg1*^floxed/+^ (*Brg1*^fl/+^) *Actin*-*creER* mouse embryonic stem cells, we performed single cell gel electrophoresis (comet assay) and observed that the deletion of a single copy of the ATPase subunit results in an increase in both medium and high amounts of DNA breaks (**Figure 5D**). This increase in DNA breaks with depleted levels of *Brg1* may explain the hightened synergistic effect observed in the presence ATRi (**Figure 5E**).

**Figure 5:**
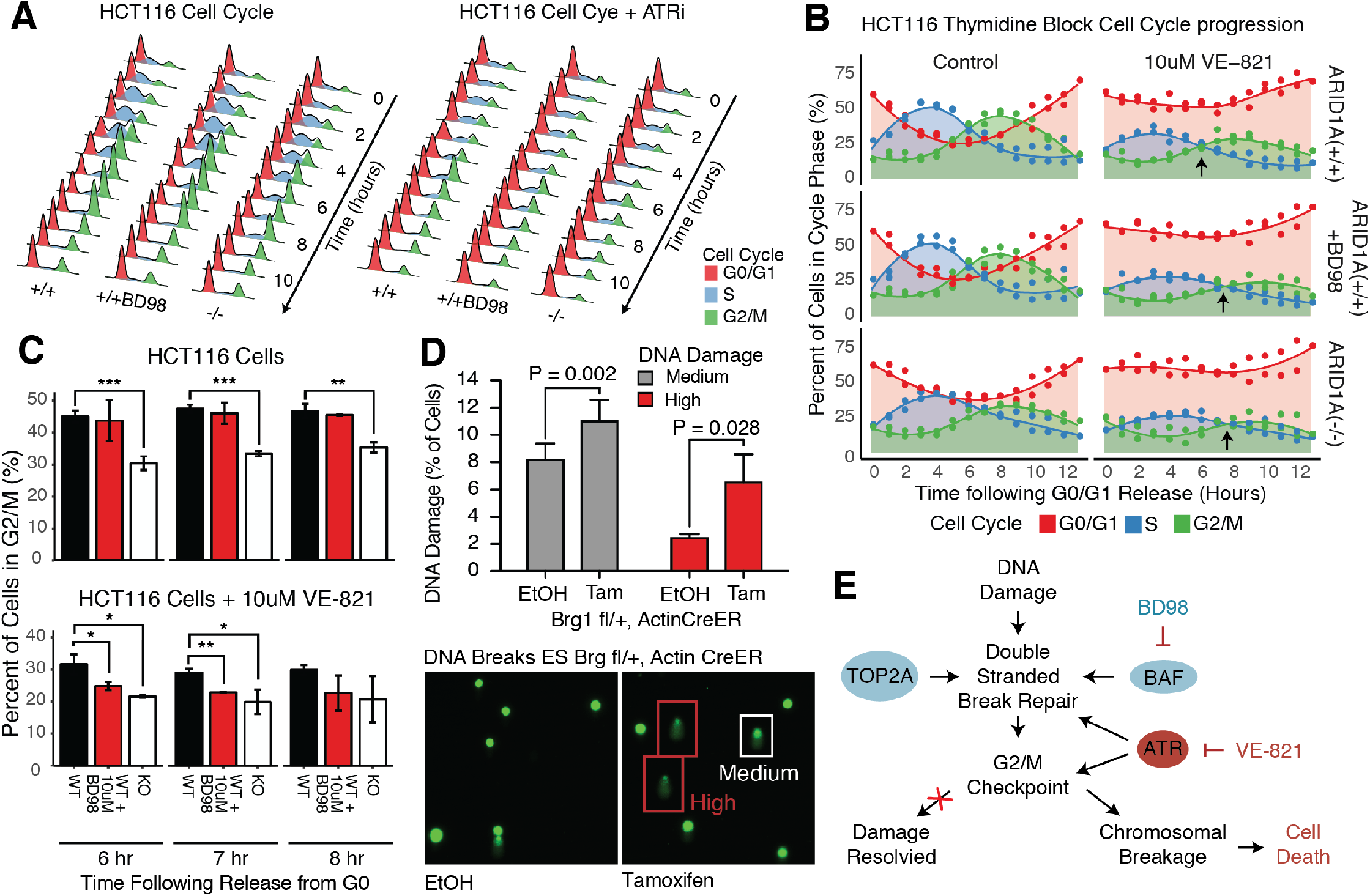
BAF inhibition results in cell cycle defects and is exacerbated by DNA damage. **(A)** Representative histograms of the cellular Propidium iodide-stained DNA content determined by FACS, in HCT116 ARID1A +/+, ARID1A −/−, and ARID1A+/+ treated with 10uM of BD98 at the indicated time points following release from cell synchronization in G0/G1, in the absence (left) or presence of 10uM VE-821 (right). Time points begin at maximum G0/G1 of second division following release from thymidine block (10hr post release). **(B)** Time course illustrating the percentage of cells in G0/G1 (red), S (blue), or G2/M (green) phase of synchronously growing HCT116 cells (as treated in 5A). **(C)** Percent differences of ARID1A null or ED89 treated cells in G2 following 6, 7, or 8 hours post-release from G0, as compared to wildtype. **(D)** Comet analyses of *Brg1*^floxed/+^ (*Brg1*^fl/+^) *Actin*-*creER m*ESCs. (Left) Quantitative analysis of three independent experiments showing the percentage of cells with medium or high levels of DNA breaks. (Right) Representative images.**(E)** Model of BD98/ATRi induced hyper-synthetic lethality mechanism and sensitivity to DNA damage response.

### SWI/SNF inhibitor effects are exacerbated by DNA damage

Recent studies have indicated a role of SWI/SNF remodeling complexes in DNA damage repair through either ATR/ATM-dependent phosphorylation of BAF170, which recruits BAF to double stranded breaks (DSB) repair sites^15,43^ or through resolving DNA decatenation through a direct interaction with TOP2A^13^. Previously, it was reported that knockdown of TOP2A in HCT116 *ARID1A*^−/−^ cells resulted in high levels of cell death, which was presumed to be a result of a stronger dependency on TOP2A upon loss of ARID1A^34^. To assess the mechanism of ATR-dependent BAF sensitivity in cancer cells and whether it was truly TOP2A-dependent, synergy was assessed between VE-821 and TOP2A/B inhibitors in WT and KO ARID1A HCT116 cells. Both cell lines were treated with increasing doses of the TOP2A inhibitor Etoposide, and the TOP2B inhibitor (XK469)^44^ for 5 days **(Supplemental Figure 2A)** with increasing doses of VE-821 **(Figure 6A**, **Supplemental Figure 2B)**, and the synergy between them was calculated (**Figure 6B-C**, **Supplemental Figure 2C**). Modest synergy was observed when both wild type and ARID1A knockout cells were treated with the TOP2B inhibitor (CI_XK469(+/+)_ = 0.75±0.05, CI_XK469(−/−)_ = 0.85±0.09) (**Supplemental Figure 2C**). However, VE-821 and Etoposide demonstrated moderate synergy in the *ARID1A*^+/+^, but “strong synergy” was observed in the knockout (**Figure 6B-C**). To confirm the added synergism observed upon functional inhibition of ARID1A, the assay was repeated with both knockdown of ARID1A, and treatment with 10μM BD98. In all three cases of deletion, knockdown, and chemical inhibition, loss of ARID1A functionality resulted in statistically stronger responses to co-administration of TOP2A and ATR inhibition. This suggests that in the absence of functional canonical BAF complexes, by either chemical or genetic deletion, cancer cells may be unable to repair DSBs which results in rapid cell death following checkpoint bypass due to ATR inhibition.

**Figure 6:**
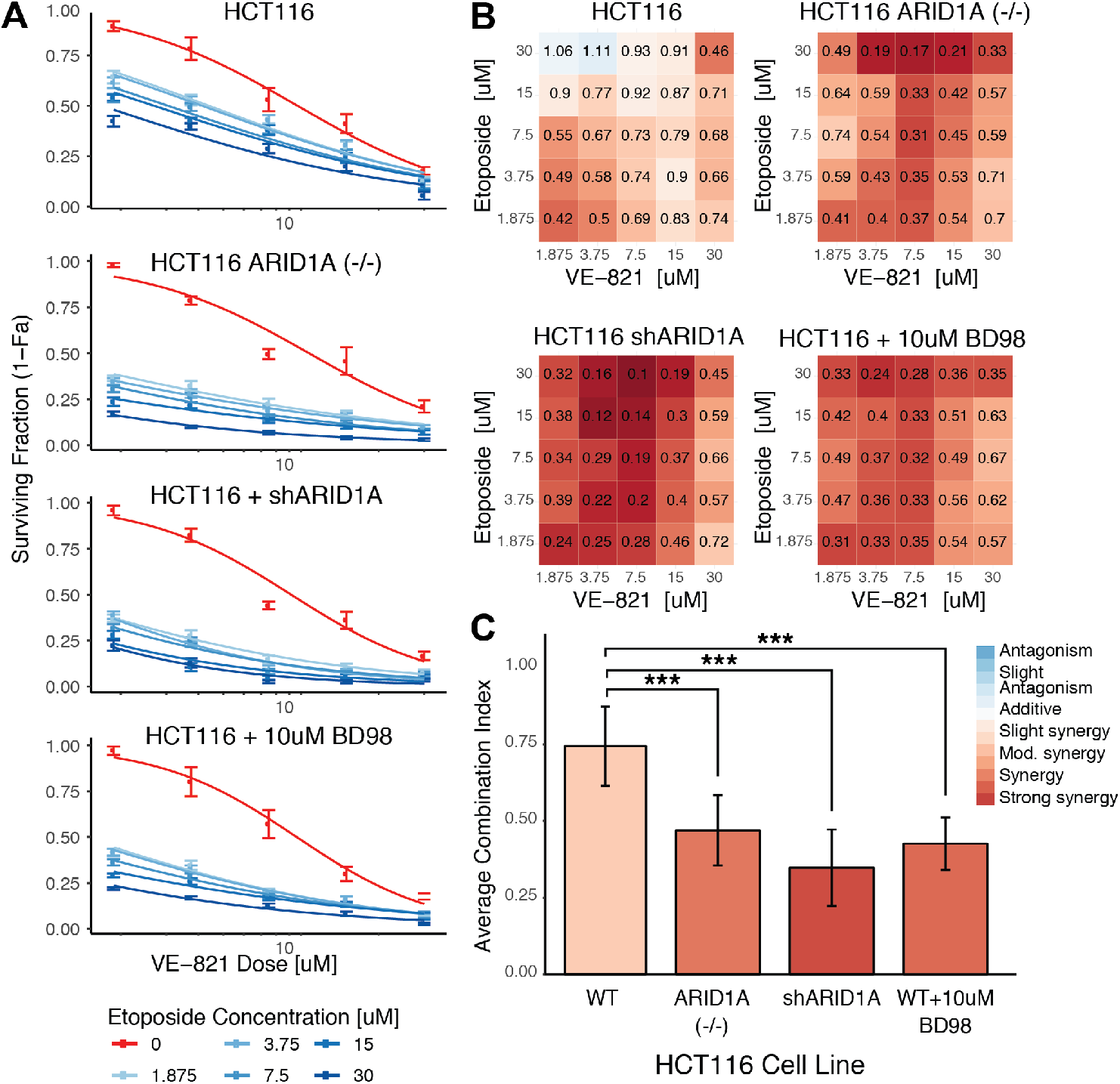
Sensitivity of ARID1A deficient cells to VE-821 and Etoposide. **(A)** Shifting dose responses of HCT116 cells, ARID1A−/− HCT116, shARID1A cells, and HCT116 cells treated with BD98, following 5-days of treatment with 25 dose combinations of VE-821 and Etoposide. **(B)** Combination index grids of cell lines (as treated in 6A). **(C)** Average combination indexes for HCT116 treated with Etoposide and VE-821 (as treated in 6A).

### BAF inhibition in mutant SWI/SNF cell lines

In ARID1A-deficient tumors, ATR inhibition is thought to trigger premature mitotic entry and chromosome instability that cannot be resolved, resulting in mitotic catastrophe^34^. Recent reports have demonstrated that loss-of-function BRG1 mutations have increased sensitivity to topoisomerase II inhibition in the presence of PRC2 inhibitors^45^, and that cancers with oncogenic BAF mutations may also exhibit increased sensitivity to PRC2 inhibition^46^. We sought to assess which cancers, including BAF-deficient cancers, might be susceptible to ATR sensitization by inhibiting SWI/SNF pathways. We tested the efficacy of BD98 and ATRi on six additional human cancer cell lines including two highly mutated proliferative cells lines (Cal-51 and MCF-7), two synovial sarcoma cell lines (Aska, Yamato) which are driven by non-canonical BAF functions (ncBAF), and two renal-cell carcinoma lines (A-704, ACHN), which harbor a mutation in the PBAF-specific subunit, *PBRM1*^39^ or are heterozygous mutants for PBRM1, respectively. Interestingly, Cal-51 is ARID1A null **(Figure S3A)**, but appears to be ARID1B-dependent, as we were unable to successfully knock down ARID1B with well-established knockdown constructs **(Figure S3B)**^47^.

To assess synergy, each cell line was treated with combinations of 5 doses of VE-821 and 5 doses of BD98 as described above. Both synovial sarcoma cell lines demonstrated no measurable dose-response to BD98 **(Figure 7A-B)**, and no quantifiable synergy between BD98 and VE-821 could be obtained. This supports the evidence that BD98 is likely acting on a canonical BAF pathway, as it has recently been discovered that the proliferation of synovial sarcoma and malignant rhabdoid tumors are dependent on ncBAF complexes, which contain neither ARID1A or ARID1B, among other subunits **(Figure 1A)**^28^. Interestingly, BD98 sensitized both MCF-7 and Cal-51 cells lines to ATR inhibition (CI_MCF-7_ = 0.45±0.04, CI_Cal-51_ = .57±0.09) (**Figure 7C-E**), despite the total loss of ARID1A in the Cal 51 line. Because Cal-51 cells have bi-allelic loss of *ARID1A* **(Supplemental Figure 3A)**, the synergy observed in this line most likely results from the actions of the redundant ARID1B^4,48^, but highlights the non-obvious nature of identifying potential clinical target populations. Finally, in both renal-cell carcinoma lines (ACHN, A-704), the drug combination demonstrated no measurable synergy (**Supplemental Figure 3C-D**). Together, these results suggest that BD98’s mechanism is acting through an ARID1-related pathway, as synovial sarcoma’s have no canonical BAF and the dependency of renal clear cell carcinomas proliferation is on PBAF.

**Figure 7:**
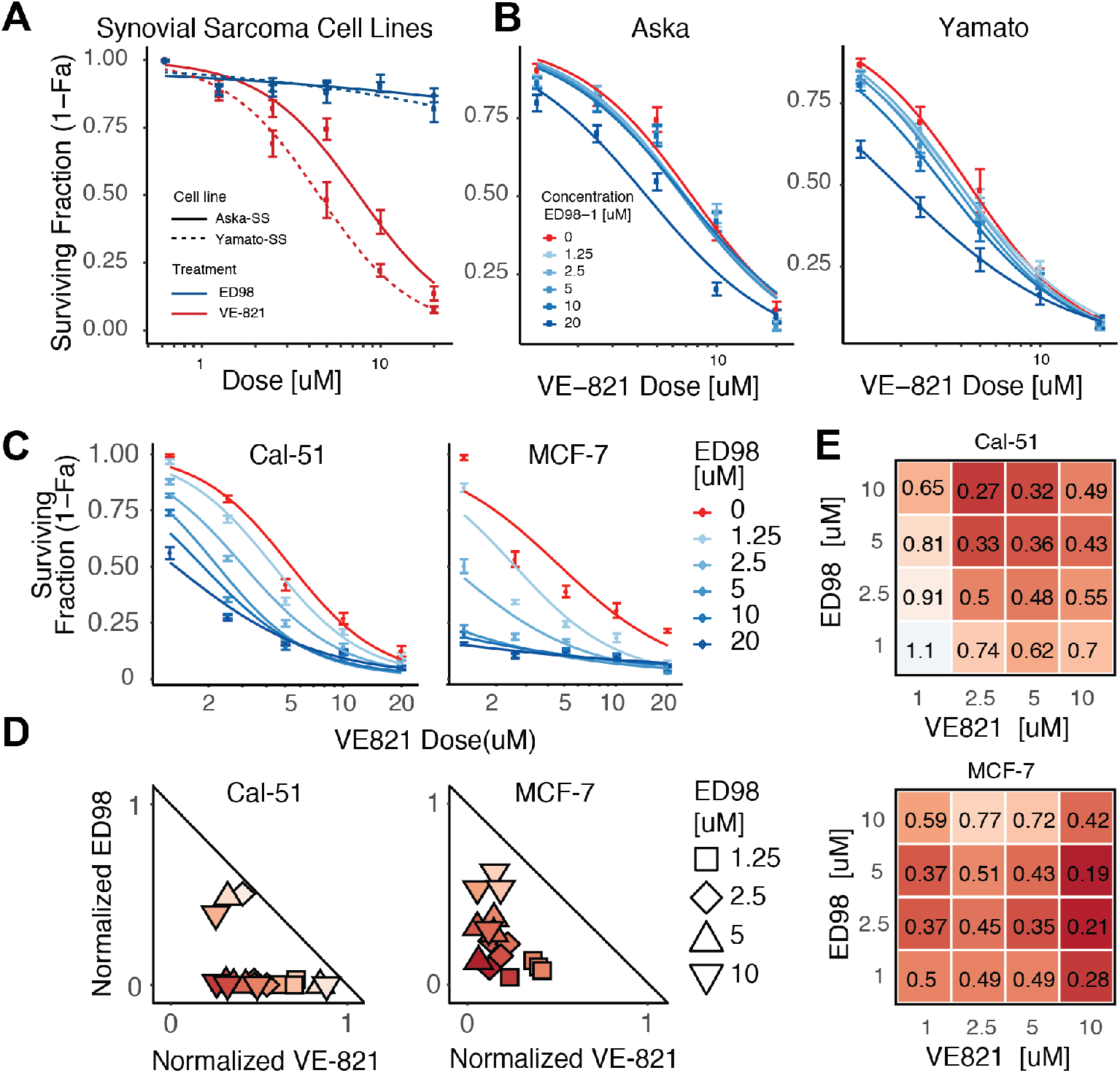
BD98 sensitization in SWI/SNF mutant cell lines. **(A)** Dose responses of synovial sarcoma cell lines treated with VE-821 or ED89. **(B)** No observable shifting dose response curves of synovial sarcoma cell lines treated with increasing combinations of VE-821 and ED89. **(C)** Shifting dose response curves of Cal-51 and MCF7 cell lines treated with increasing combinations of VE-821 and ED89. **(C)** Normalized isobolograms of Cal-51 and MCF7 cell lines treated with increasing combinations of VE-821 and ED89. **(C)** Combination index plots of Cal-51 and MCF7 cell lines treated with increasing combinations of VE-821 and BD98.

## Discussion

In this study, we assess whether small-molecule inhibition of a specialized function of canonical BAF complexes can serve as a viable therapeutic strategy for the treatment of cancer. The 15 subunits of BAF complexes are combinatorially assembled from the products of 31 genes giving remarkable biologic specificity to these complexes^49,50^. This conceptual advance led us to search for inhibitors of the specific repressive function of canonical BAF complexes in the hope of avoiding toxicity, while still targeting a potentially druggable aspect of these large assemblies. Previous work indicated that ATR-induced BAF phosphorylation is essential for localization to sites of DNA damage and that cancer cell lines deficient in BAF complexes exhibit synthetic lethality with a variety of clinical cancer therapeutics^34,45,51^. These observations led us to test the possibility that an inhibitor of a specialized function of BAF complexes might synergize with ATR inhibitors, allowing them to be used at lower, less toxic concentrations in the treatment of cancer. General inhibitors of BRG1/BRM ATPases have been described^21,22^ but because the loss of BRG1 is lethal in nearly all cell types, they are likely far too toxic for therapeutic use. In addition, an inhibitor of the bromodomain of BRG1/SMARCA4 has been described, but is not effective at inhibiting cancer cell growth^20^. By combining BD98 with inhibitors of the ATR kinase, we have demonstrated that cancer cells undergo a chemically-induced synthetic lethal effect, particularly in cancers whose oncogenesis is not primarily SWI/SNF-dependent. This suggests that inhibitors of canonical BAF have the potential for broader therapeutic utility, beyond classically disrupted BAF-related cancers such as synovial sarcomas^52^, malignant rhabdoid tumors^53,54^, and renal clear cell carcinomas^39,41^.

Therapeutic targeting of multi-subunit complexes has been a challenge due to the varied functions, compositions, and specificity of such complexes. Suspected inhibitors of protein complexes likely lack substrate affinity for a single subunit, but rather interact with composite surfaces produced by combinatorial assembly. This poses significant barriers to traditional biochemical characterizations, as demonstrated by our prior efforts to identify the specific target of interaction^27^. Further, challenges surrounding complex purification, along with the lack of structural insights into such complexes (as there is no published crystal structure) has kept the development of specific inhibitors largely out-of-reach. Based on previous work^15,34^, we reasoned that small molecules that block canonical ARID1A-containing BAF complexes may also be synergistic with ATR inhibition. Many of the putative BAF inhibitors that we discovered in our primary screens were highly toxic, as would be expected based on the essential role of most subunits of the BAF complex. However, specific molecules that block the transcriptional repressive function of BAF complexes appear non-toxic and yet we show are highly synergistic with ATR inhibition in several human cell lines. Of the initial hits that arose from our previous screen^24,27^, we have observed that the macrolactam candidates exhibited exquisite stereospecifity which will aid in the development of better and more potent inhibitors of SWI/SNF-dependent pathways. Despite the observation of strong synergy between BD98 and ATRi, challenges remain to transition these compounds into animal models. These macrocyclic inhibitors are synthesized through a nucleophilic aromatic substitution intramolecular ring-forming process with stereochemical dependencies^55^, and as such, scale-up remains a bottleneck and the limiting quantities of compound can be prohibitive to downstream characterizations. This further highlights the need for early validation assays that require very little compound, such as the synergy method employed in this work (Chory & Divakaran, unpublished).

Our studies and previous observations indicate that the mechanism underlying synergy between BD98 and ATR inhibition is likely related to ATR/ATM phosphorylation of BRG1 and BAF170 promoting the interaction of BAF with sites of DNA damage^15^, which is further exacerabated upon treatment with inhibitors of TOP2A **(Figure 5E)**. Therefore, we hypothesize that synergy of the two inhibitors arrises from the roles of ATR in recruiting BAF to sites of DNA damage, BAF-dependent DNA damage repair, and role of ATR in checkpoint arrest. Failure at each of these contingent steps could lead to synergistic cell death. While acute ARID1A loss has been shown to provide synthetic lethality with ATR inhibition^34^, the role of ATR in the localization of BAF to sites of DNA damage suggest a broader role of BD98, and like molecules, in treating cancer than just cancers that have mutations in ARID1A or ARID1B. This, coupled with recent findings that tumor cells deficient in BAF complexes exhibit synthetic lethality from a variety of chemotherapeutics (anti-PD-1, checkpoint kinase inhibitors, EZH2 inhibitors)^34,45,51^, suggests that chemical inhibition of BAF complexes may have synergistic value by sensitizing cancers to a range of therapeutics.

## METHODS

### Cell culture

HCT116 ARID1A(+/+) and ARID1A(−/−) cell lines were courtesy of Dr. Diana Hargreaves, and originally obtained from Horizon Discovery. Cancer cell lines were obtained from American Type Tissue Collection. All cancer cell lines were grown in McCoy’s 5a Media supplemented with 10% FBS. mES cells were cultured with standard conditions on gelatin coated plates in DMEM media containing 7.5% FBS, 7.5% Serum-replacement and LIF. Media was replenished daily, and cells were passaged every 48 hours. For complete reagents, cell line details, and methods see Supplemental Experimental Procedures.

### Synergy viability assays

Viability assays were performed in 384-well plates. Cells were plated at 500 cells/well. 25 dose combinations were administered 24 hours after seeding and media containing fresh drug was replaced every 48 hours. After 5 days, cellular viability was measured and combination index values were calculated utilizing R (R-package: EChormatin/SynergyCalc”), as described by Chou and Talalay^35^. See Supplemental Experimental Procedures.

### Lentiviral preparation and infection

Lentiviruses were produced LentiX-HEK293T cells via spinfection with polyethylenimine transfection. HEK293T cells were cultured using standard conditions in DMEM media containing 10% FBS. HEK293T cells were transfected with PEI with lentiviral pLKO shARID1A, GIPZ shARID2, GIPZ shARID1B knockdown vectors, co-transfected with packaging vectors psPAX2 and pMD2.G as previously described^56^. Viral supernatant was collected and used to for spin-fection. Infected cells were selected with puromycin 48h following transfection and maintained under selection for 2-3 passages. See Supplemental Experimental Procedures.

### RT-qPCR analysis

RNA was extracted from cells; cDNA was synthesized, and samples were analyzed by qPCR. See Supplemental Experimental Procedures and Supplemental Table 1.

### Cell synchronization

HC T116 cells were plated 6-well plates and incubated with thymidine for 18hr, released into fresh media for 8 h, and re-incubated with thymidine again for 16hr, washed several times with PBS, and released into fresh media to synchronize into G1/early S and then collected at respective time points. The mESCs were incubated with thymidine for 8h, released into fresh media for 7 h, and then incubated with thymidine again for 7h. Cells were washed several times with PBS, released into fresh media, and collected at respective time points. See Supplemental Experimental Procedures.

### Cell cycle analysis

For hour-by-hour analysis, cancer cells were collected, rinsed with PBS, fixed in 70% ethanol. Cells were fixed overnight, pelleted, rinsed in PBS, stained with Propidium Iodide, and analyzed by flow cytometry. For mESCs, cell cycle analysis was performed by staining with BrdU-FITC/7-AAD and analyzed by flow cytometry. See Supplemental Experimental Procedures.

### Comet Assay

mES cells were dissociated and analyzed with single cell gel electrophoresis. After staining with vista Green DNA dye, comet images were captured by fluorescence microscopy. See Supplemental Experimental Procedures.

### Western Blot

Nuclear extracts were prepared by benzonase treatment and nuclear extract buffer containing HEPES, KCl, EDTA, MgCl2, Glycerol, NP-40 Protease inhibitors, DTT, and Benzonase (See supplemental experimental procedures for full details. Nuclei were lysed in RIPA buffer and protein concentrations were determined by Bradford assay (Biorad). Proteins were separated by SDS–PAGE electrophoresis with a 4–12% Bis-Tris protein gel (Thermo Scientific) and then transferred to an Immobilon-FL membrane (Millipore). Blots were probed with primary antibodies, rabbit α-Arid1a(D2A8u) (1:1000, Cell Signalling #12354), and mouse α-Arid1b (D2D) (1:1000, Novus Biologicals # H00057492-M01) followed by fluorescence-conjugated secondary antibodies (Li-Cor) and bands were detected using an Odyssey CLX imaging system (Li-Cor).

## ACKNOWLEDGEMENTS

We thank S. Divakaran and D. Hargreaves for helpful discussions. HCT116 *ARID1A(−/−)* cells were a gift from D. Hargreaves (Salk Institute). D. Hargreaves provided ARID1B knockdown constructs. JR Raab (UNC) provided ARID1A knockdown constructs. The Aska-SS and Yamato-SS cell lines were provided by Kazuyuki Itoh, Norifume Naka, and Satoshi Takenaka (Osaka University, Japan). E.J.C. was partially supported by an NSF Graduate Research Fellowship and the Ruth L. Kirschstein National Research Service Award (F31 CA203228-02). J.G.K. was partially supported by PHS Grant Number T32 CA09151, awarded by the National Cancer Institute, DHHS. The initial screen and follow-up studies of top inhibitors were supported by NIH grant DA032469 and the Showalter Trust. E.C.D. is supported by The V Foundation for Cancer Research (V2014-004 and D2016-030) and the NIH (CA207532). G.R.C. is supported by NIH grants (CA163915, NS046789), CIRM RB4-05886, SFARI and the Howard Hughes Medical Institute.

## Author Contributions

E.J.C designed and conducted the experiments, analyzed data, and wrote the manuscript. J.G.K., C.Y.C, S.G., and V.D.D. designed and conducted experiments. G.R.C and E.C.D. designed experiments and wrote the manuscript.

## Supplemental Information

**Figure S1:**
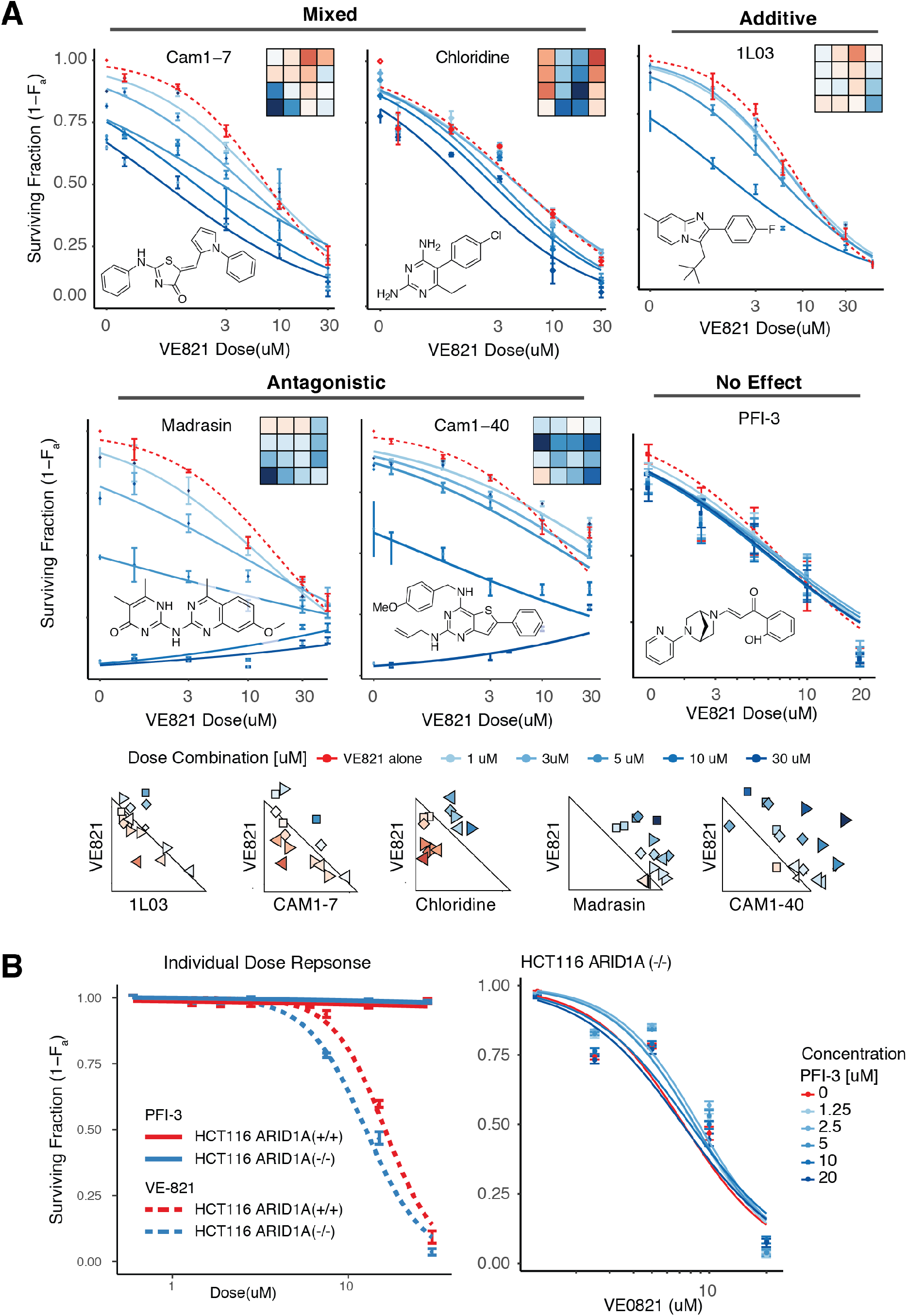
Synergy analysis amoung putative SWI/SNF inhibitors. (A) Shifting dose response curves and synergy analysis of 6 putative SWI/SNF inhibitors. (B) Left: Dose response curves of HCT116 cell line exposed to increasing concentrations of ATR inhibitor (VE-821) and PFI-3 for 5 days. Right: Shifting IC50 dose-response curves of HCT116 cells treated with VE-821 and increasing doses of PFI-3 for 5 days.

**Figure S2:**
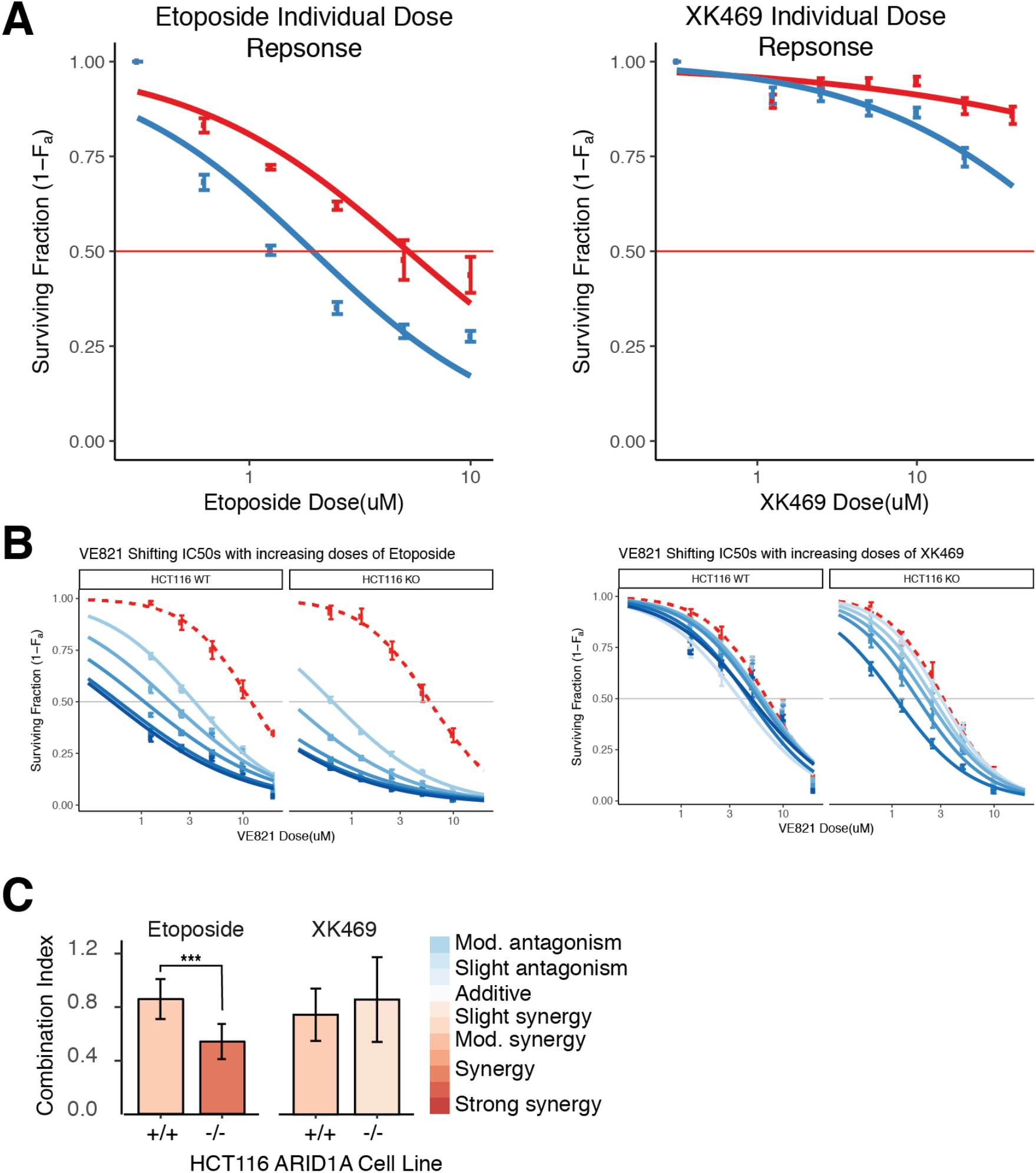
Dose response curves and shifting IC50 plots of Topoisomerase inhibitors. **(A)** Dose response curves of wild type HCT116 cells (Blue) and ARID1A −/− HCT116 cells (red) exposed to increasing concentrations of ATR inhibitor (VE-821) and topoisomerase inhibitors (Etoposide and XK469) for 5 days. **(B)** Shifting IC50 curves wild type HCT116 cells and ARID1A −/− HCT116 cells exposed to increasing concentrations of ATR inhibitor (VE-821) and topoisomerase inhibitors (Etoposide and XK469) for 5 days**. (C)** Synergy assessment of HCT116 cells and ARID1A −/− HCT116 cells exposed to increasing concentrations of ATR inhibitor (VE-821) and topoisomerase inhibitors (Etoposide and XK469) for 5 days.

**Figure S3:**
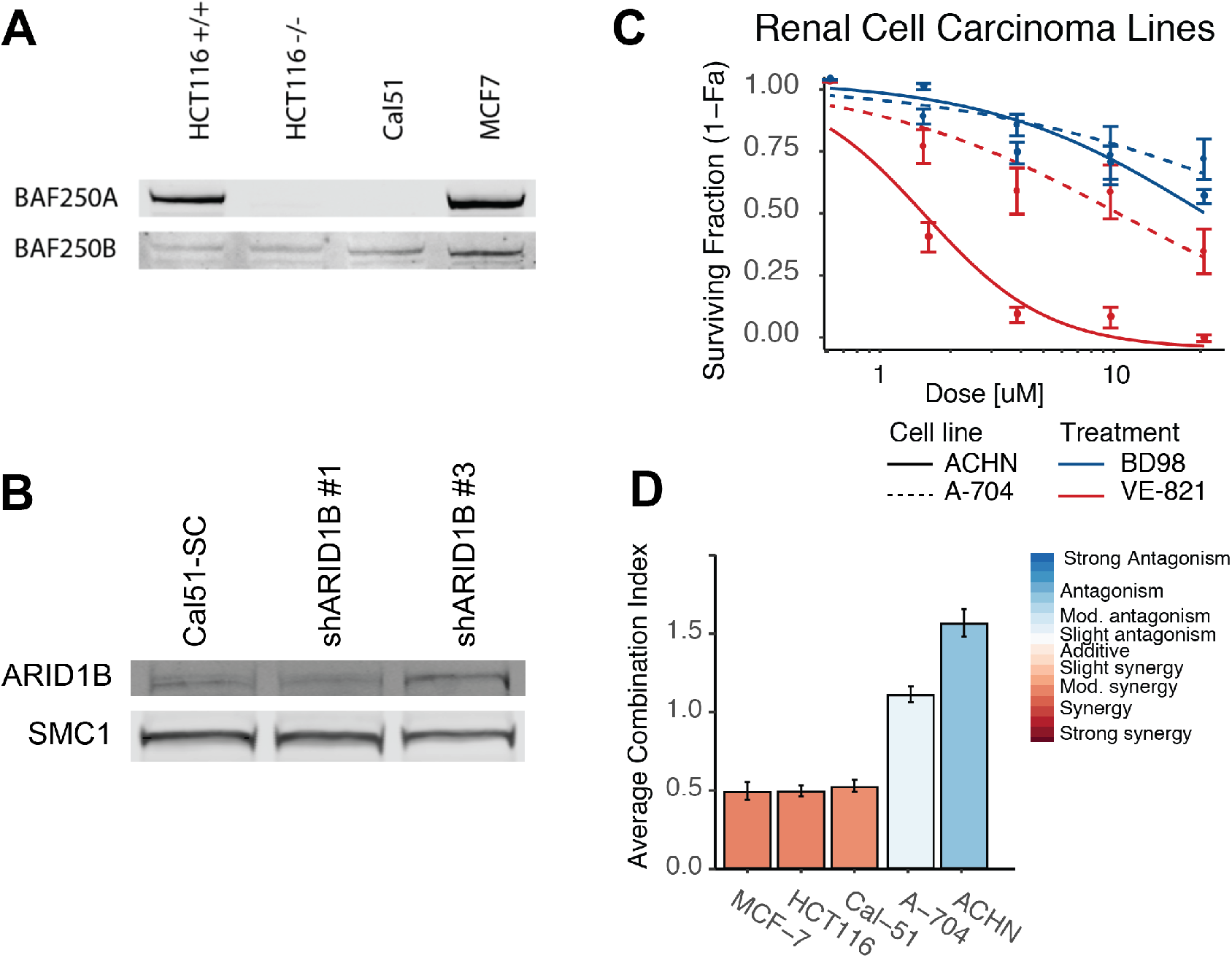
ARID1A/b Mutant Cancer lines. **(A)** Expression of ARID1A and ARID1B in HCT116, ARID1A −/− HCT116, Cal51 and MCF7 cancer cell lines. **(B)** Attempted knockdown of ARID1B in Cal51 cell lines. **(C)** Dose response of Renal cell carcinoma lines to increasing concentrations of ATR inhibitor (VE-821) and BD98. **(D)** Synergy assessment in SWI/SNF mutant cancer cell lines.

**Table S1.**
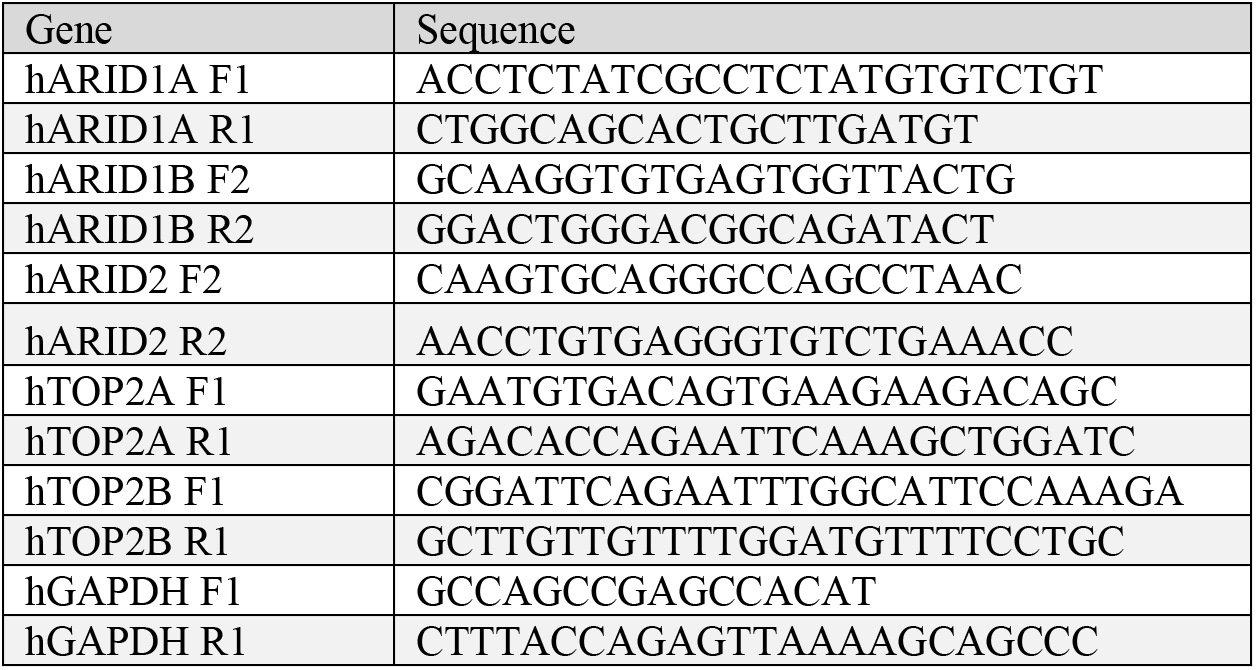
qPCR primers.

## Supplemental Experimental Procedures

### Cell culture

Isogenic HCT116 ARID1A(+/+) and ARID1A(−/−) cell lines were courtesy of Dr. Diana Hargreaves, and originally obtained from Horizon Discovery. The Aska-SS and Yamato-SS cell lines were provided by Kazuyuki Itoh, Norifume Naka, and Satoshi Takenaka (Osaka University, Japan). MCF-7, Cal-51, A-704, and ACHN cell lines were obtained from American Type Tissue Collection. All cancer cell lines were grown in McCoy’s 5a Media supplemented with 10% FBS. mES cells were cultured on gelatin coated plates in DMEM media (Life Technologies) containing 7.5% ES-sure FBS (Applied StemCell), 7.5% KnockOut SR (Life Technologies), HEPES buffer (Life Technologies), Glutamax (Life Technologies), Penicillin-Streptomycin (Life Technologies), 2-mercapto-ethanol (Life Technologies), and MEM-NEAA (Life Technologies), and LIF. Media was replenished daily and cells were passaged every 48 hours.

### Chemicals

VE-821 was purchased from Selleck Chem. BD98 was synthesized as described by Fitzgerald and colleagues^57^. Chloridine (Pyrimethamine), Madrasin, Etoposide, and XK469 were purchased from Sigma-Aldrich. Cam1-7 and Cam1-40 (Pubchem CID 49792165) were purchased through Evotec. 1L03 (Pubchem CID 46902783), 2A03 (Pubchem CID 54631898), and 2C05 (Pubchem CID 54631408) were provided by the Broad Institute.

### Synergy viability assays

Viability assays were performed in 384-well plates. Cells were plated at 500 cells/well. Each drug was administered in 5-doses with 4 replicates per plate, and each 5×5 drug combination was administered with 8 replicates per plate, 24 hours after seeding. All error bars represent 4 or 8 technical replicates averaged over 3 independent experiments. Media containing fresh drug was replaced every 48 hours. After 5 days, cellular viability was measured using CellTitre blue, as described by the manufacturer. Combination index values were calculated utilizing R (R-package: EChormatin/SynergyCalc”), as described by Chou and Talalay^35^.

### Lentiviral preparation and infection

Lentiviruses were produced HEK 293 cells (ATCC) via spinfection with polyethylenimine transfection. HEK 293 cells (ATCC) were cultured using standard conditions in DMEM media (Life Technologies) containing 10% FBS (Applied StemCell). and Penicillin-Streptomycin (Life Technologies). HEK 293 cells were transfected with PEI (Polysciences Inc., 24765) with lentiviral pLKO shARID1A, GIPZ shARID2, or GIPZ shARID1B knockdown vectors, co-transfected with packaging vectors psPAX2 and pMD2.G as previously described^56^. PLKO shRNA construct targeting ARID1A (TRCN0000059090: shRNA-2) was a gift from Dr. Jesse R. Raab^58^, GIPZ Human ARID2 shRNA was purchased from Dharmacon (CloneId:V2LHS_74399), and GIPZ shARID1B was provided by Dr. Diana Hargreaves^47^ (originally from Dharmacon). 12h after transfection, media was changed, after another 48h, media was collected and supernatant was used to spinfect cells in the presence of 10 μg/ml Polybrene (Santa Cruz Biotechnology) at 1000g for 1hr. Infected cells were selected with 2 μg/ml puromycin beginning 48hr after infection, and maintained under selection for 2-3 passages.

### RT-qPCR analysis

RNA was extracted from cells using Trisure (Bioline) and cDNA was synthesized from 1ug RNA using the SensiFAST SYBR Lo-Rox (Bioline). Delta Samples were run on a QuantStudio 6 Flex system (Life Technologies). 2^−ΔΔCT^ was calculated as described by Livak and Schmittgen ^59^ where: −ΔΔCT = (CT_GOI_ − CT_Gapdh_)*ARID1A*(−/−) − (CT_GOI_ −CT_Gapdh_)*ARID1A*(+/+). Primers for qPCR are included in Supplementary Table 1.

### Cell synchronization

HCT116 cells were plated at 2E6 cells/well in 6-well plates and incubated with 2mM thymidine for 18hr, released into fresh media for 8 h, and incubated with thymidine again for 16hr, washed several times with PBS, and released into fresh media containing either 10uM BD98, 10uM VE-821, or both to synchronize into G1/early S. To begin collection at the maximum percent of cells in G0/G1, cells were allowed to proceed through one cell cycle for 9 hours, and then collected at respective time points. The maximum G0/G1 time point occurred at 10 hours post-release from thymidine block. The mESCs were incubated with 2mM thymidine for 7–8h, released into fresh media for 7 h, and then incubated with thymidine again for 7h. Cells were washed several times with PBS, released into fresh media, and collected at respective time points.

### Cell cycle analysis

For hour-by-hour analysis, cells were collected, rinsed with PBS, and vortexed while adding 1mL ice cold 70% ethanol. Cells were fixed overnight, pelleted at 1000xg, rinsed in PBS, resuspended in PBS containing 50 ug/mL RnaseA and 10 ug/mL Propidium Iodide and incubated at 37C for 30min. Flow cytometry analysis was performed on a BD Accuri Flow Cytometer. Individual cells were gated based on forward and side scatter, auto-fluorescent cells were omitted, and remaining cells were then analyzed for propidium iodide levels. To determine the percent of cells in each phase, DNA content histograms histograms were analyzed using R-package “mixtools” EM algorithm for mixtures of univariate normals. The areas of each mixed normal distribution were calculated to represent the total number of cells in each phase of the cell cycle. For mESCs, cell cycle analysis was performed using BD Biosciences BrdU-FITC FACS kits. The mESCs were incubated with BrdU for 1h, stained with 7-AAD, and analyzed on a BD FACScan.

### Comet Assay

Briefly, the mES cells were dissociated by 0.05% trypsin, and 1,000 cells were used in single cell gel electrophoresis (comet assay) with the Cell BioLab’s OxiSelect Comet Assay Kit according to the manufacturer’s instructions. Alkaline solution was used for electrophoresis with 15V, ~270-300mA for 22 mins. After staining with vista Green DNA dye, comet images were captured by fluorescence microscopy. For each experiment, >300 cells were counted from >20 images per experiment for 3-4 experiments for each condition. In each sample, the percentage of cells with medium and high tail moments was calculated to represent the cells with DNA breaks. *Brg1*^floxed/+^ *Actin*-*creER* mESCs were treated with EtOH for 1uM tamoxifen for 24h, and passaged 3-5 times before the analysis.

## REFERENCES

(1) Peterson, C. L.; Herskowitz, I. Characterization of the Yeast SWI1, SWI2, and SWI3 Genes, Which Encode a Global Activator of Transcription. Cell 68 (3), 573–583. https://doi.org/10.1016/0092-8674(92)90192-F.

(2) Khavari, P.; Peterson, C.; Tamkun, J.; Mendel, D. BRG 1 Contains a Conserved Domain of the SWI 2/SNF 2 Family Necessary for Normal Mitotic Growth and Transcription. Nature 1993.

(3) Hodges, C.; Kirkland, J. G.; Crabtree, G. R. The Many Roles of BAF (MSWI/SNF) and PBAF Complexes in Cancer. Cold Spring Harbor perspectives in medicine 6 (8), a026930. https://doi.org/10.1101/cshperspect.a026930.

(4) Mathur, R.; Alver, B. H.; Roman, A. K.; Wilson, B. G.; Wang, X.; Agoston, A. T.; Park, P. J.; Shivdasani, R. A.; Roberts, C. W. ARID1A Loss Impairs Enhancer-Mediated Gene Regulation and Drives Colon Cancer in Mice. Nature Genetics 49 (2), 296–302. https://doi.org/10.1038/ng.3744.

(5) Lessard, J.; Wu, J. I.; Ranish, J. A.; Wan, M.; Winslow, M. M.; Staahl, B. T.; Wu, H.; Aebersold, R.; Graef, I. A.; Crabtree, G. R. An Essential Switch in Subunit Composition of a Chromatin Remodeling Complex during Neural Development. Neuron 2007, 55 (2), 201–215. https://doi.org/10.1016/j.neuron.2007.06.019.

(6) Meisenberg, C.; Downs, J. A. The SWI/SNF Chromatin Remodelling Complex: Its Role in Maintaining Genome Stability and Preventing Tumourigenesis. DNA Repair 32, 127–133. https://doi.org/10.1016/j.dnarep.2015.04.023.

(7) Rafati, H.; Parra, M.; Hakre, S.; Moshkin, Y.; Verdin, E.; Mahmoudi, T. Repressive LTR Nucleosome Positioning by the BAF Complex Is Required for HIV Latency. PLoS Biology 9 (11), e1001206–20. https://doi.org/10.1371/journal.pbio.1001206.

(8) Mahmoudi, T.; Parra, M.; Vries, R. G.; Kauder, S. E.; Verrijzer, P. C.; Ott, M.; Verdin, E. The SWI/SNF Chromatin-Remodeling Complex Is a Cofactor for Tat Transactivation of the HIV Promoter. Journal of Biological Chemistry 281 (29), 19960–19968. https://doi.org/10.1074/jbc.M603336200.

(9) Conrad, R. J.; Fozouni, P.; Thomas, S.; Sy, H.; Zhang, Q.; Zhou, M.-M.; Ott, M. The Short Isoform of BRD4 Promotes HIV-1 Latency by Engaging Repressive SWI/SNF Chromatin-Remodeling Complexes. Molecular Cell. https://doi.org/10.1016/j.molcel.2017.07.025.

(10) Wang, X.; Wang, S.; Troisi, E. C.; Howard, T. P.; Haswell, J. R.; Wolf, B. K.; Hawk, W. H.; Ramos, P.; Oberlick, E. M.; Tzvetkov, E. P.; et al. BRD9 Defines a SWI/SNF Sub-Complex and Constitutes a Specific Vulnerability in Malignant Rhabdoid Tumors. Nat Commun 2019, 10 (1), 1881. https://doi.org/10.1038/s41467-019-09891-7.

(11) Brien, G. L.; Remillard, D.; Shi, J.; Hemming, M. L.; Chabon, J.; Wynne, K.; Dillon, E. T.; Cagney, G.; Mierlo, G.; Baltissen, M. P.; et al. Targeted Degradation of BRD9 Reverses Oncogenic Gene Expression in Synovial Sarcoma. Elife 2018, 7, e41305. https://doi.org/10.7554/elife.41305.

(12) Ho, L.; Ronan, J. L.; Wu, J.; Staahl, B. T.; Chen, L.; Kuo, A.; Lessard, J.; Nesvizhskii, A. I.; Ranish, J.; Crabtree, G. R. An Embryonic Stem Cell Chromatin Remodeling Complex, EsBAF, Is Essential for Embryonic Stem Cell Self-Renewal and Pluripotency. Proceedings of the National Academy of Sciences 106 (13), 5181–5186. https://doi.org/10.1073/pnas.0812889106.

(13) Dykhuizen, E. C.; Hargreaves, D. C.; Miller, E. L.; Cui, K.; Korshunov, A.; Kool, M.; Pfister, S.; Cho, Y.-J.; Zhao, K.; Crabtree, G. R. BAF Complexes Facilitate Decatenation of DNA by Topoisomerase IIα. Nature 497 (7451), 624–627. https://doi.org/10.1038/nature12146.

(14) Miller, E. L.; Hargreaves, D. C.; Kadoch, C.; Chang, C.-Y.; Calarco, J. P.; Hodges, C.; Buenrostro, J. D.; Cui, K.; Greenleaf, W. J.; Zhao, K.; et al. TOP2 Synergizes with BAF Chromatin Remodeling for Both Resolution and Formation of Facultative Heterochromatin. Nature Structural & Molecular Biology 33, 1492. https://doi.org/10.1038/nsmb.3384.

(15) Peng, G.; Yim, E.-K.; Dai, H.; Jackson, A. P.; van der Burgt, I.; Pan, M.-R.; Hu, R.; Li, K.; Lin, S.-Y. BRIT1/MCPH1 Links Chromatin Remodelling to DNA Damage Response. Nature cell biology 11 (7), 865–872. https://doi.org/10.1038/ncb1895.

(16) Wang, X.; Lee, R. S.; Alver, B. H.; Haswell, J. R.; Wang, S.; Mieczkowski, J.; Drier, Y.; Gillespie, S. M.; Archer, T. C.; Wu, J. N.; et al. SMARCB1-Mediated SWI/SNF Complex Function Is Essential for Enhancer Regulation. Nature Genetics 49 (2), 289–295. https://doi.org/10.1038/ng.3746.

(17) Yoo, A. S.; Staahl, B. T.; Chen, L.; Crabtree, G. R. MicroRNA-Mediated Switching of Chromatin-Remodelling Complexes in Neural Development. Nature 460 (7255), 642–646. https://doi.org/10.1038/nature08139.

(18) Kadoch, C.; Hargreaves, D. C.; Hodges, C.; Elias, L.; Ho, L.; Ranish, J.; Crabtree, G. R. Proteomic and Bioinformatic Analysis of Mammalian SWI/SNF Complexes Identifies Extensive Roles in Human Malignancy. Nature Genetics 45 (6), 592–601. https://doi.org/10.1038/ng.2628.

(19) Fedorov, O.; Castex, J.; Tallant, C.; Owen, D.; Martin, S.; Aldeghi, M.; Monteiro, O.; Filippakopoulos, P.; Picaud, S.; Trzupek, J.; et al. Selective Targeting of the BRG/PB1 Bromodomains Impairs Embryonic and Trophoblast Stem Cell Maintenance. Science Advances 1(10), e1500723–e1500723. https://doi.org/10.1126/sciadv.1500723.

(20) Vangamudi, B.; Paul, T. A.; Shah, P. K.; Kost-Alimova, M.; Nottebaum, L.; Shi, X.; Zhan, Y.; Leo, E.; Mahadeshwar, H. S.; Protopopov, A.; et al. The SMARCA2/4 ATPase Domain Surpasses the Bromodomain as a Drug Target in SWI/SNF-Mutant Cancers: Insights from CDNA Rescue and PFI-3 Inhibitor Studies. Cancer Research 75 (18), 3865–3878. https://doi.org/10.1158/0008-5472.CAN-14-3798.

(21) Muthuswami, R.; Mesner, L.; Wang, D.; Hill, D.; Imbalzano, A.; Hocken-smith, J. Phosphoaminoglycosides Inhibit SWI2/SNF2 Family DNA-Dependent Molecular Motor Domains. Biochemistry 39 (15), 4358–4365. https://doi.org/10.1021/bi992503r.

(22) Papillon, J. P.; Nakajima, K.; Adair, C. D.; Hempel, J.; Jouk, A. O.; Karki, R. G.; Mathieu, S.; Möbitz, H.; Ntaganda, R.; Smith, T.; et al. Discovery of Orally Active Inhibitors of Brahma Homolog (BRM)/SMARCA2 ATPase Activity for the Treatment of Brahma Related Gene 1 (BRG1)/SMARCA4-Mutant Cancers. Journal of medicinal chemistry 2018, 61 (22), 10155–10172. https://doi.org/10.1021/acs.jmed-chem.8b01318.

(23) Ho, L.; Miller, E. L.; Ronan, J. L.; Ho, W.; Jothi, R.; Crabtree, G. R. EsBAF Facilitates Pluripotency by Conditioning the Genome for LIF/STAT3 Signalling and by Regulating Polycomb Function. Nature cell biology 13 (8), 903–913. https://doi.org/10.1038/ncb2285.

(24) Dykhuizen, E.; Carmody, L.; Tolliday, N.; Crabtree, G.; Palmer, M. Screening for Inhibitors of an Essential Chromatin Remodeler in Mouse Embryonic Stem Cells by Monitoring Transcriptional Regulation. Journal of Biomolecular Screening 17 (9), 1221–1230. https://doi.org/10.1177/1087057112455060.

(25) Tan, D. Diversity-Oriented Synthesis: Exploring the Intersections between Chemistry and Biology. Nature Chemical Biology 2005, 1 (2), 74–84. https://doi.org/10.1038/nchembio0705-74.

(26) National Center for Biotechnology Information. Pubchem BioAssay Database; AID=602436, Source=Broad Institute. 2012.

(27) Marian, C. A.; Stoszko, M.; Wang, L.; Leighty, M. W.; de Crignis, E.; Maschinot, C. A.; Gatchalian, J.; Carter, B. C.; Chowdhury, B.; Hargreaves, D. C.; et al. Small Molecule Targeting of Specific BAF (MSWI/ SNF) Complexes for HIV Latency Reversal. Cell Chem Biol 2018. https://doi.org/10.1016/j.chembiol.2018.08.004.

(28) Michel, B. C.; D’Avino, A. R.; Cassel, S. H.; Mashtalir, N.; McKenzie, Z. M.; McBride, M. J.; Valencia, A. M.; Zhou, Q.; Bocker, M.; Mares, L.; et al. A Non-Canonical SWI/SNF Complex Is a Synthetic Lethal Target in Cancers Driven by BAF Complex Perturbation. Nature Cell Biology 2018, 20 (12), 1410–1420. https://doi.org/10.1038/s41556-018-0221-1.

(29) Shiloh, Y.; Ziv, Y. The ATM Protein Kinase: Regulating the Cellular Response to Genotoxic Stress, and More. Nature Reviews Molecular Cell Biology 14 (4), 197–210. https://doi.org/10.1038/nrm3546.

(30) Zou, L.; Elledge, S. J. Sensing DNA Damage through ATRIP Recognition of RPA-SsDNA Complexes. Science 300 (5625), 1542–1548. https://doi.org/10.1126/science.1083430.

(31) Stoszko, M.; Crignis, E.; Rokx, C.; Khalid, M.; Lungu, C.; Palstra, R.; Kan, T.; Boucher, C.; Verbon, A.; Dykhuizen, E. C.; et al. Small Molecule Inhibitors of BAF; A Promising Family of Compounds in HIV-1 Latency Reversal. EBioMedicine 3, 108–121. https://doi.org/10.1016/j.ebiom.2015.11.047.

(32) Karnitz, L. M.; Zou, L. Molecular Pathways: Targeting ATR in Cancer Therapy. Clinical cancer research : an official journal of the American Association for Cancer Research 21 (21), 4780–4785. https://doi.org/10.1158/1078-0432.CCR-15-0479.

(33) Weber, A.; Ryan, A. ATM and ATR as Therapeutic Targets in Cancer. Pharmacology and Therapeutics 149 (C), 124–138. https://doi.org/10.1016/j.pharmthera.2014.12.001.

(34) Williamson, C. T.; Miller, R.; Pemberton, H. N.; Jones, S. E.; Campbell, J.; Konde, A.; Badham, N.; Rafiq, R.; Brough, R.; Gulati, A.; et al. ATR Inhibitors as a Synthetic Lethal Therapy for Tumours Deficient in ARID1A. Nature Communications 7, 13837. https://doi.org/10.1038/ncomms13837.

(35) Chou, T. Theoretical Basis, Experimental Design, and Computerized Simulation of Synergism and Antagonism in Drug Combination Studies. Pharmacological Reviews 58 (3), 621–681. https://doi.org/10.1124/pr.58.3.10.

(36) Chou, T. Drug Combination Studies and Their Synergy Quantification Using the Chou-Talalay Method. Cancer Research 70 (2), 440–446. https://doi.org/10.1158/0008-5472.CAN-09-1947.

(37) Chou, T.; Talalay, P. Quantitative Analysis of Dose-Effect Relationships: The Combined Effects of Multiple Drugs or Enzyme Inhibitors. Advances in enzyme regulation 1984, 22, 27–55. https://doi.org/10.1016/0065-2571(84)90007-4.

(38) Brownlee, P. M.; Chambers, A. L.; Cloney, R.; Bianchi, A.; Downs, J. A. BAF180 Promotes Cohesion and Prevents Genome Instability and Aneuploidy. CellReports 6 (6), 973–981. https://doi.org/10.1016/j.celrep.2014.02.012.

(39) Chowdhury, B.; Porter, E. G.; Stewart, J. C.; Ferreira, C. R.; Schipma, M. J.; Dykhuizen, E. C. PBRM1 Regulates the Expression of Genes Involved in Metabolism and Cell Adhesion in Renal Clear Cell Carcinoma. PLoS ONE 2016, 11 (4), e0153718. https://doi.org/10.1371/journal.pone.0153718.

(40) Jiao, Y.; Pawlik, T. M.; Anders, R. A.; Selaru, F. M.; Streppel, M. M.; Lucas, D. J.; Niknafs, N.; Guthrie, V.; Maitra, A.; Argani, P.; et al. Exome Sequencing Identifies Frequent Inactivating Mutations in BAP1, ARID1A and PBRM1 in Intrahepatic Cholangiocarcinomas. Nature Genetics 45 (12), 1470–1473. https://doi.org/10.1038/ng.2813.

(41) Goeppinger, S.; Keith, M.; Tagscherer, K. E.; Singer, S.; Winkler, J.; Hoffmann, T. G.; Pahernik, S.; Duensing, S.; Hohenfellner, M.; Kopitz, J.; et al. PBRM1 (BAF180) Protein Is Functionally Regulated by P53‐induced Protein Degradation in Renal Cell Carcinomas. The Journal of Pathology 237 (4), 460–471. https://doi.org/10.1002/path.4592.

(42) Takebayashi, S.-I.; Lei, I.; Ryba, T.; Sasaki, T.; Dileep, V.; Battaglia, D.; Gao, X.; Fang, P.; Fan, Y.; Esteban, M. A.; et al. Murine EsBAF Chromatin Remodeling Complex Subunits BAF250a and Brg1 Are Necessary to Maintain and Reprogram Pluripotency-Specific Replication Timing of Select Replication Domains. Epigenetics & Chromatin 6 (1), 42. https://doi.org/10.1186/1756-8935-6-42.

(43) Kwon, S.-J.; Park, J.-H.; Park, E.-J.; Lee, S.-A.; Lee, H.-S.; Kang, S.; Kwon, J. ATM-Mediated Phosphorylation of the Chromatin Remodeling Enzyme BRG1 Modulates DNA Double-Strand Break Repair. Oncogene 34 (3), 303–313. https://doi.org/10.1038/onc.2013.556.

(44) Gao, H.; Huang, K.; Yamasaki, E.; Chan, K.; Chohan, L.; Snapka, R. XK469, a Selective Topoisomerase IIbeta Poison. Proceedings of the National Academy of Sciences 96 (21), 12168–12173.

(45) Fillmore, C. M.; Xu, C.; Desai, P. T.; Berry, J. M.; Rowbotham, S. P.; Lin, Y.-J.; Zhang, H.; Marquez, V. E.; Hammerman, P. S.; Wong, K.-K.; et al. EZH2 Inhibition Sensitizes BRG1 and EGFR Mutant Lung Tumours to TopoII Inhibitors. Nature 520 (7546), 239–242. https://doi.org/10.1038/nature14122.

(46) Pang, B.; de Jong, J.; Qiao, X.; Wessels, L. F.; Neefjes, J. Chemical Profiling of the Genome with Anti-Cancer Drugs Defines Target Specificities. Nature Chemical Biology 11 (7), 472–480. https://doi.org/10.1038/nchembio.1811.

(47) Kelso, T. W.; Porter, D. K.; Amaral, M. L.; Shokhirev, M. N.; Benner, C.; Hargreaves, D. C. Chromatin Accessibility Underlies Synthetic Lethality of SWI/SNF Subunits in ARID1A-Mutant Cancers. eLife 2017, 6. https://doi.org/10.7554/eLife.30506.

(48) Helming, K. C.; Wang, X.; Wilson, B. G.; Vazquez, F.; Haswell, J. R.; Manchester, H. E.; Kim, Y.; Kryukov, G. V.; Ghandi, M.; Aguirre, A. J.; et al. ARID1B Is a Specific Vulnerability in ARID1A-Mutant Cancers. Nat Med 2014, 20 (3), 251–254. https://doi.org/10.1038/nm.3480.

(49) Wu, J. I.; Lessard, J.; Crabtree, G. R. Understanding the Words of Chromatin Regulation. Cell 2009, 136 (2), 200–206. https://doi.org/10.1016/j.cell.2009.01.009.

(50) Kadoch, C.; Crabtree, G. R. Mammalian SWI/SNF Chromatin Remodeling Complexes and Cancer: Mechanistic Insights Gained from Human Genomics. Science Advances 1 (5), e1500447–e1500447. https://doi.org/10.1126/sciadv.1500447.

(51) Pan, D.; Kobayashi, A.; Jiang, P.; de Andrade, L.; Tay, R.; Luoma, A.; Tsoucas, D.; Qiu, X.; Lim, K.; Rao, P.; et al. A Major Chromatin Regulator Determines Resistance of Tumor Cells to T Cell–Mediated Killing. Science 359 (6377), eaao1710–775. https://doi.org/10.1126/science.aao1710.

(52) Kadoch, C.; Crabtree, G. R. Reversible Disruption of MSWI/SNF (BAF) Complexes by the SS18-SSX Oncogenic Fusion in Synovial Sarcoma. Cell 2013, 153 (1), 71–85. https://doi.org/10.1016/j.cell.2013.02.036.

(53) Roberts, C. W.; Galusha, S. A.; McMenamin, M. E.; Fletcher, C. D.; Orkin, S. H. Haploinsufficiency of Snf5 (Integrase Interactor 1) Predisposes to Malignant Rhabdoid Tumors in Mice. Proc National Acad Sci 2000, 97 (25), 13796–13800. https://doi.org/10.1073/pnas.250492697.

(54) Wilson, B. G.; Wang, X.; Shen, X.; McKenna, E. S.; Lemieux, M. E.; Cho, Y.-J.; Koellhoffer, E. C.; Pomeroy, S. L.; Orkin, S. H.; Roberts, C. W. Epigenetic Antagonism between Polycomb and SWI/SNF Complexes during Oncogenic Transformation. Cancer Cell 18 (4), 316–328. https://doi.org/10.1016/j.ccr.2010.09.006.

(55) Marcaurelle, L. A.; Comer, E.; Dandapani, S.; Duvall, J. R.; Gerard, B.; Kesavan, S.; Maurice, L. D.; Liu, H.; Lowe, J. T.; Marie, J.-C.; et al. An Aldol-Based Build/Couple/Pair Strategy for the Synthesis of Medium- and Large-Sized Rings: Discovery of Macrocyclic Histone Deacetylase Inhibitors. Journal of the American Chemical … 132 (47), 16962–16976. https://doi.org/10.1021/ja105119r.

(56) Tiscornia, G.; Singer, O.; Verma, I. M. Design and Cloning of Lentiviral Vectors Expressing Small Interfering RNAs. Nature Protocols 2006, 1 (1), 234–240. https://doi.org/10.1038/nprot.2006.36.

(57) Fitzgerald, M. E.; Mulrooney, C. A.; Duvall, J. R.; Wei, J.; Suh, B.-C.; Akella, L. B.; Vrcic, A.; Marcaurelle, L. A. Build/Couple/Pair Strategy for the Synthesis of Stereochemically Diverse Macrolactams via Head-to-Tail Cyclization. ACS Combinatorial Science 14 (2), 89–96. https://doi.org/10.1021/co200161z.

(58) Raab, J. R.; Resnick, S.; Magnuson, T. Genome-Wide Transcriptional Regulation Mediated by Biochemically Distinct SWI/SNF Complexes. PLoS genetics 11 (12), e1005748. https://doi.org/10.1371/journal.pgen.1005748.

(59) Schmittgen, T. D.; Livak, K. J. Analyzing Real-Time PCRDatabythe Comparative CT Method. Nature Protocols 3 (6), 1101–1108. https://doi.org/10.1038/nprot.2008.73.

